# TCR-engineered T-cells directed against Ropporin-1 constitute a safe and effective treatment for triple-negative breast cancer in near-clinical models

**DOI:** 10.1101/2024.01.10.574978

**Authors:** Dian Kortleve, Dora Hammerl, Mandy v Brakel, Rebecca Wijers, Daphne Roelofs, Kim Kroese, Mieke Timmermans, Chen-Yi Liao, Anita Trapman-Jansen, Renée Foekens, Justine Michaux, Monique de Beijer, Sonja I. Buschow, Jeroen A.A. Demmers, Marleen Kok, Erik H.J. Danen, Michal Bassani-Sternberg, John W. Martens, Rachel J.M. Abbott, Reno Debets

## Abstract

Triple-negative breast cancer (TNBC) shows an urgent need for new therapies. We discovered Ropporin-1 (ROPN1) as a target to treat TNBC with T-cells. ROPN1 showed high and homogenous expression in 90% of primary and metastatic TNBC but not in healthy tissues. HLA-A2-binding peptides were detected via immunopeptidomics and predictions and used to retrieve T-cell receptors (TCRs) from naïve repertoires. Following gene introduction into T-cells and stringent selection, we retrieved a highly specific TCR directed against the epitope FLYTYIAKV that did not recognize non-cognate epitopes from alternative source proteins. Notably, this TCR mediated killing of three-dimensional tumoroids *in vitro* and tumor cells *in vivo* and outperformed standard-of-care drugs. Finally, the T-cell product expressing this TCR and manufactured using a clinical protocol fulfilled standard safety and efficacy assays. Collectively, we have identified and preclinically validated ROPN1 as a target and anti-ROPN1 TCR T-cells as a treatment for the vast majority of TNBC patients.

---

Triple-negative breast cancer (TNBC) is an aggressive subtype of breast cancer (BC) that affects relatively young women at an age of 40-50 years at diagnosis and accounts for 15-20% of all BC cases. The 5-year overall survival rate for metastatic TNBC is only 12%^1^ which reflects the lack of effective treatment options. Indeed, TNBC lacks targets for hormonal therapy and HER2 amplification and patients with TNBC are therefore not eligible for standard therapies for BC^2^. The poly ADP-ribose polymerase (PARP) inhibitors olaparib and talazoparib have shown effectiveness in treating advanced TNBC with germline Breast Cancer gene (BRCA) mutations (overall response rate (ORR) 55-62%)^3,4^, yet only 10-20% of TNBC patients carry these mutations^5–7^ and the relatively high ORR doesn’t translate into overall survival benefit^8^. The Programmed Cell Death 1 (PD-1) inhibitor pembrolizumab, when combined with chemotherapy, improves responses for a subgroup of metastatic TNBC, being positive for Programmed Cell Death Ligand 1 (PD-L1), resulting in an ORR of 53% compared to 41% for chemotherapy only, but this subgroup only accounts for 30-40% of all patients with TNBC^9^. A more recently approved standard-of-care option for TNBC patients is the Trophoblast cell surface antigen 2 (TROP2)-targeting antibody-drug conjugate Sacituzumab-govitecan, which has resulted in an ORR of 35% and showed a gain in ORR of 30% when compared to chemotherapy in a cohort of relapsed TNBC patients. Treatment with Sacituzumab-govitecan, however, is accompanied with serious side effects with >50% of patients experiencing adverse events of grade 3 or higher, such as neutropenia and leukopenia^10,11^. Despite new drugs having become available, there is still an unmet demand for effective and safe therapies for TNBC patients.

A promising new treatment option for TNBC is adoptive T-cell therapy (ACT). ACT is a form of immune therapy that relies on the transfer of tumor-specific T-cells that are generally generated by inserting a gene to express either a chimeric antigen receptor (CAR) or T-cell receptor (TCR) into the patients’ T-cells *ex vivo*. CAR-T-cells constitute a major breakthrough for the treatment for different B cell malignancies^12–16^ as well as multiple myeloma^17–19^ with six of these CAR T-cell products now implemented as standard-of-care^20^. Unfortunately, the efficacy of CAR T-cells to treat solid tumors lags behind significantly to that of hematological tumors^21–26^. Explanations may include that in general solid tumors lack tumor-specific targets available on the surface of solid tumors and harbor an immunosuppressive tumor microenvironment^27^.

TCRs recognize targets through presentation via major histocompatibility complex (MHC) molecules, not being restricted to cell surface antigens and broadening the spectrum of potential tumor target antigens. Notably, TCR-engineered T-cells targeting either intracellular New York Esophageal Squamous Cell Carcinoma 1 (NY-ESO1) or MAGE family member A4 (MAGE-A4) in the context of human leukocyte antigen (HLA-)A2 have shown clear clinical responses with ORRs up to 61% in patient with melanoma and synovial sarcoma. Both TCR T-cell products have good safety profiles and the latter is even close to market approval^28–30^. However, high expression of NY-ESO-1 and MAGE-A4 is limited to above-mentioned solid tumor types, and expressions are often heterogeneous and variable among patients^31^.

In the current study we have identified and tested new target antigens and corresponding TCRs in a stepwise manner and according to stringent selection criteria for safety and efficacy to enable the development of ACT for TNBC. In our search, only targets were included that were absent in healthy tissues to reduce the risk of on-target toxicity. At the same time target selection included high and homogenous expression in TNBC to facilitate effective anti-tumor T-cell responsiveness^32^. TCRs were retrieved from naïve repertoires and selected when harboring a stringent epitope-recognition motif to reduce the risk of off-target toxicity^33–35^ <u>and</u> driving eradication of TNBC in advanced tumor models. Overall, our work reports the discovery of a novel target, the Cancer Germline Antigen (CGA) Ropporin-1 (ROPN1), as well as the retrieval and extensive preclinical validation of an anti-ROPN1 TCR for the treatment of the majority of TNBC patients.

## Results

### ROPN1 is absent in healthy tissues and shows abundant and homogenous expression in primary and metastatic TNBC

To discover a target suitable for ACT for TNBC, we have applied a stepwise and stringent selection process whereby large gene expression datasets of both normal healthy and tumor tissues were screened for intracellular antigens^37^ (see for details below and **Figure 1a**).

**Figure 1.**
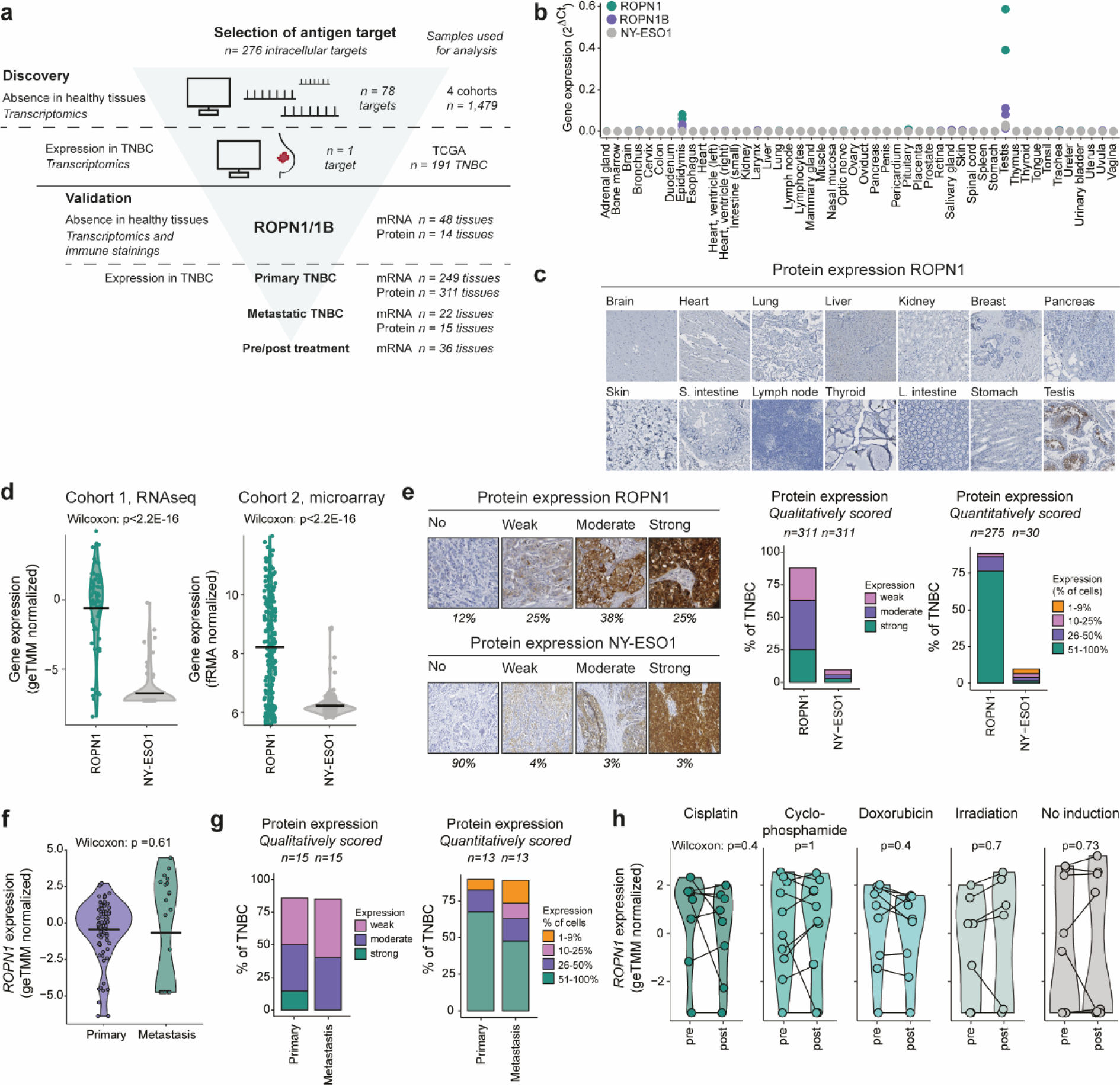
ROPN1/B is absent in healthy tissues and highly and homogenously present in primary and metastatic TNBC independent of pre-treatment. **a)** Flow chart of the discovery of ROPN1/B as a target for T-cell treatment of TNBC and validation of its tumor-specific expression using healthy tissues, as well as primary and metastatic TNBC tissues. Intracellular targets (Cancer Germline Antigens: n=276) were screened for absent expression in healthy tissues (n=1,479 tissues) and expression in TNBC (n=191 tissues). ROPN1/B mRNA and protein expression was further validated in multiple sets of healthy tissues and cohorts of patient tumor tissues. **b)** Dot plot shows gene expressions as fold change relative to GAPDH (2^-dCt^) according to qPCR using a cDNA library of 48 healthy tissue samples. NY-ESO1 was used as a reference. Green: ROPN1; purple: ROPN1B; grey: NY-ESO1 (CTAG1B), n=2-3 per target antigen. **c)** Representative immune stainings of ROPN1/B using an array of 14 healthy tissues (2-6 donors per tissue, total n=66). **d)** Violin plots show distribution of gene expression of ROPN1 (green) and NY-ESO1 (CTAG1B, grey) in TNBC cohort 1 (n=66, geTMM normalized) and cohort 2 (n=183, fRMA normalized) (see Methods for details on cohorts). Data was analyzed using the Wilcoxon signed-rank test: cohort 1: p<2.2E-16; cohort 2: p<2.2E-16. **e)** Representative immune stainings of TNBC with different intensities for ROPN1/B or NY-ESO1 (left panel). Stacked bar graphs (middle panel) show fraction of TNBC tumors with weak, moderate and strong immune staining of ROPN1/B or NY-ESO1 (tissue microarrays (TMA), n=311). Stacked bar graphs (right panel) show fraction of TNBC tumors with either 1-9%, 10-25%, 26-50% or 51-100% of tumors cells positive for ROPN1/B or NY-ESO1 protein. **f)** Violin plot shows distribution of gene expressions of ROPN1/B in primary and metastatic TNBC retrieved from TNBC cohort 1 (n=66, geTMM normalized) and cohort 3 (n=22) following batch correction. Data was analyzed using the Wilcoxon signed-rank test: p=0.61. **g)** Stacked bar graphs (left panel) show fraction of primary and metastatic TNBC with weak, moderate and strong immune staining of ROPN1/B (whole tissue sections, n=15 paired samples). Stacked bar graphs (right panel) show fraction of primary and metastatic TNBC with either 1-9%, 10-25%, 26-50% or 51-100% of tumor cells positive for ROPN1/B. **h)** Violin plots show distribution of gene expression of ROPN1/B in pre- and post-induction treatment biopsies of metastatic TNBC retrieved from cohort 4 (n=53 of which n=44 are paired samples, geTMM normalized). Data was analyzed using the paired pre-post samples with the Wilcoxon signed-rank test: Cisplatin: n=8, p=0.4; Cyclophosphamide: n=10, p=1; Doxorubicin: n=9, p=0.4; Irradiation: n=7, p=0.7; No induction: n=10, p=0.73. Abbreviations in figure: ACT: Adoptive T-cell therapy; TNBC: Triple Negative Breast Cancer; S. intestine: small intestine; L. intestine: Large intestine.

In a first step, we observed that ROPN1 as well as its paralog ROPN1B (>95% amino acid sequence homology) was one of 78 intracellular proteins that was absent in large series of healthy organs according to multiple gene expression databases (**Extended Data figure 1a**). Gene and protein expression of ROPN1 in healthy tissues was confirmed to be limited to the testis, and to a lesser extent, the epididymis (**Figure 1b-c**). Both these tissues are immune-privileged^36,37^, meaning these tissues are not seen by T-cells as they lack MHC molecules.

In a second step, the screening of tumor samples revealed that ROPN1 was expressed in >90% of patients with TNBC. In comparison, targets for ACT with clinical precedent, such as NY-ESO1 and MAGE-A4, were only expressed in 16% and 25% of TNBC patients, respectively (**Extended Data figure 1b**). Furthermore, the levels of gene and protein expression were significantly higher for ROPN1 (**Figure 1d-e, Extended Data figure 1b**) when compared to above mentioned reference targets. Strikingly, ROPN1 protein was detected in 50-100% of tumor cells in about 75% of primary TNBC (**Figure 1e**). These levels of abundance, extent of expression and homogeneity are, to our knowledge, unique when compared to other targets of ACT. To assess the robustness of ROPN1 as a target for ACT, we also investigated gene- and protein expressions in metastatic TNBC lesions (**Figure 1f-g**) and observed that the majority of studied lymph node lesions expressed ROPN1 in >50% of tumor cells, highly concordant with the extent of expression observed in primary tumors (**Figure 1g**). In addition, ROPN1 expression in distant TNBC lesions was not affected by pre-operative treatments, such as cisplatin, cyclophosphamide, doxorubicin or radiation (**Figure 1h**). Notably, ROPN1 was also expressed in 90% of skin cutaneous melanoma (SKCM) (**Extended Data figure 1c**), implying that these patient cohorts could also benefit from ACT targeting ROPN1.

### HLA-A2-restricted and endogenously processed peptides enable retrieval of ROPN1-specific TCRs from naïve repertoires

TCRs recognize small peptide fragments of the target antigen in the context of HLA molecules on the surface of cells. We set out to investigate which peptides originating from the ROPN1/B protein were recognized by TCRs, not shared with other source proteins, and restricted to the HLA-A2 subtype, the latter covering 30-50% of the EU and US population^38^. For details on peptide selection, see **Extended Data table 1** and **Methods**. ROPN1/B proteins contain 212 amino acids and cover >200 theoretical HLA-A2-restricted peptides according to the prediction algorithm NetMHCpan 4.1. All these peptides were then filtered for *in silico* qualities that are considered relevant to antigen processing and presentation, such as location of protease cleavage sites and affinities for the transporter associated with antigen processing (TAP) and binding to HLA-A2, which shortlisted the number of peptides to 20 (**Figure 2a**). This list of potential epitopes was supplemented with 17 peptides identified following mass-spectrometry of HLA class I-bound peptides from ROPN1^+^ TNBC (2 out of 17), K562ABC cells (1 out of 17) and from datasets of primary tumor samples^39,40^ (14 out of 17) (see **Extended Data table 2**, **Methods and Supplementary methods)**. Notably, 1 peptide (FLYTIAKV) that was identified in primary tumor samples was also shortlisted based on in silico predictions, yielding a total set of 36 peptides. This set of peptides was then screened for lack of similarity with other source antigens using Expitope software^41^. The peptides that were only present in ROPN1/B needed to fulfill minimal binding to HLA-A2 *in vitro,* after which 21 peptides survived that were further assessed for two parameters: the amplitude (maximal HLA-A2 binding at saturating peptide concentration) and the half-maximal effective concentration (EC50) (for details, see **Methods**). Eleven peptides showed binding to HLA-A2 in a dose-dependent manner and were ranked based on amplitude (**Figure 2b**).

**Figure 2.**
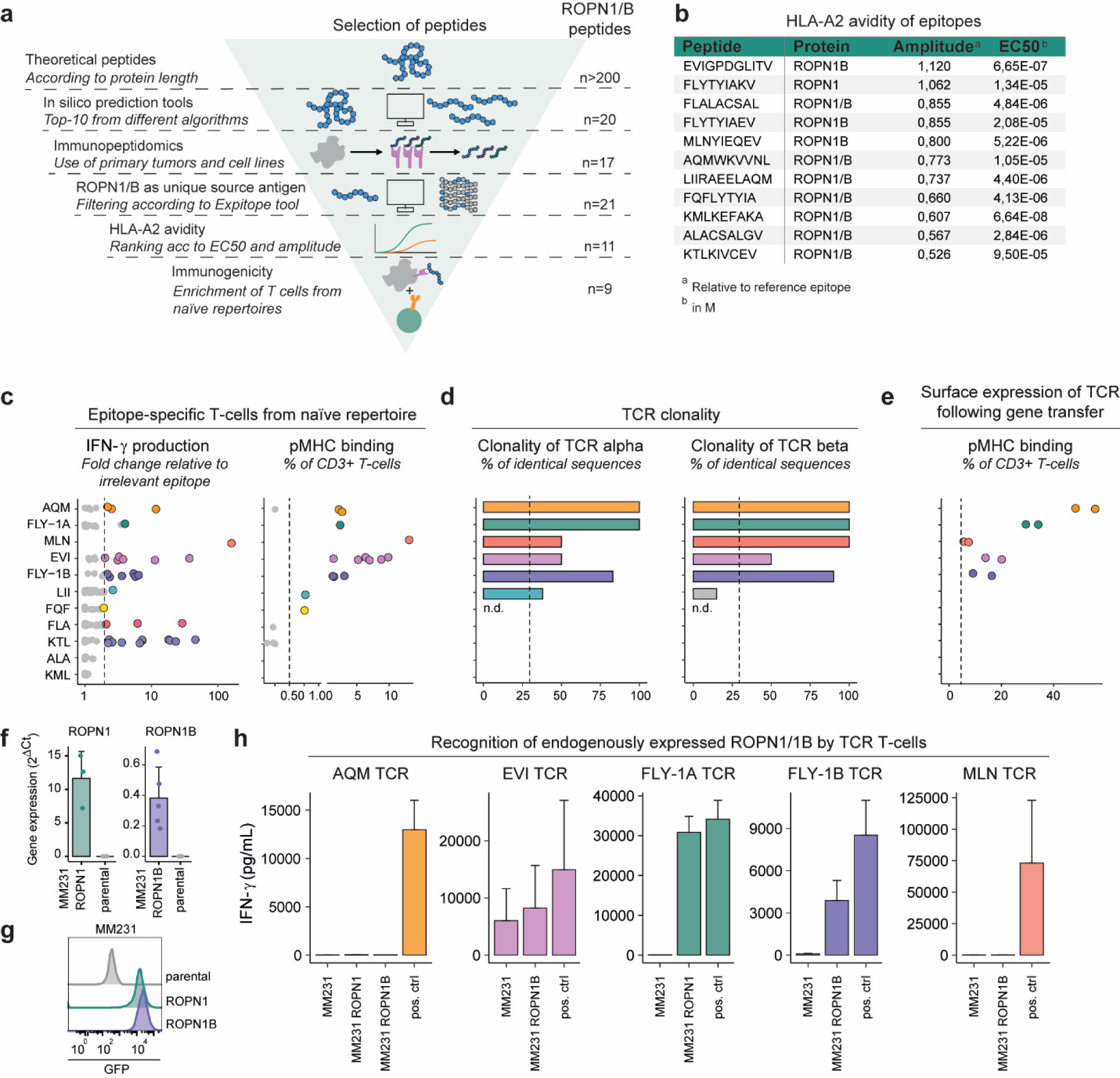
Retrieval and selection of ROPN1/B peptides according to uniqueness and HLA-A2-avidity, as well as natural TCRs against endogenously processed epitopes. **a)** Flow chart of retrieval and selection of ROPN1/B peptides according to *in silico* predictions, peptide elution, as well as non-cross-reactivity (n=21) and avidity for HLA-A2 (n=11) (see **Methods** for details on each tool/assay, and **Extended Data table 1** for an overview of the results per tool). **b)** Table presents overview of HLA-A2 binding characteristics for the shortlisted peptides that fed into T-cell enrichments. The gp100 epitope YLEPGPVTA (YLE) was used as a reference. Amplitudes are expressed as fold change (FC) of median fluorescent intensity (MFI) of anti-HLA-A2-PE relative to YLE at highest peptide concentrations, and EC50 values (mean, calculated via Graphpad Prism 5.0) are listed in Molarity (M). **c)** Dots represent enriched T-cell populations (ea. dot is one successful enrichment) according to FC in IFN-γ production compared to irrelevant epitope (RP) (left panel). Epitope stimulated T-cells which produced minimally 200 pg/mL and 2x more IFN-γ than RP stimulated T-cells were stained for pMHC binding, which is shown as percentage of CD3^+^ T-cells (right panel). **d)** Clonality of TCRα and TCRβ sequences from epitope-specific T-cell populations (from c) is shown as percentage of total number of sequences. **e)** TCRαβ combinations derived from clonal populations (from d) were introduced into T-cells and tested for TCR surface expression, which is shown as percentage of CD3^+^ T-cells. Each epitope in **c-e** is shown with a unique color and when dots are presented in grey, these epitopes didn’t meet the selection criteria for further characterization (see **Methods** and **Extended Data figure 5** for details). **f)** Bar plots represent gene expressions depicted as FC relative to GAPDH according to qPCR of ROPN1/B using ROPN1/B-GFP overexpressing MM231 cells. ROPN1 expressing MM231 cells are visualized in green (n=3), ROPN1B expressing MM231 cells in purple (n=5), and parental MM231 cells (not expressing ROPN1/B) in grey (n=5). **g)** Representative histogram shows MFI of GFP expression in ROPN1 expressing MM231 (green), ROPN1B (expressing MM231 (purple) and in parental MM231 (not expressing GFP, grey). **h)** IFN-γ production upon stimulation with the ROPN1/B expressing or parental MM231 is shown. TCR T-cells directed against ROPN1/B epitopes that showed surface TCR expression in **e** were assessed for sensitivity towards the endogenously processed and presented cognate epitope using the MM231 cell lines from **f**. Positive controls are BSM cells loaded with cognate epitope.

To retrieve ROPN1-specific T-cells, these 11peptides were individually fed to CD11c^+^ dendritic cells that were co-cultured with autologous naïve CD8^+^ T-cells from peripheral blood mononuclear cells (PBMCs) derived from a set of 13 healthy donors. Nine out of 11 peptides yielded T-cells that demonstrated significant epitope-specific IFN-γ secretion and peptide-MHC-binding (**Figure 2c, Extended Data figure 2a**). These epitope-specific T-cell populations were FACS-sorted according to peptide-MHC-binding or peptide-induced CD137 expression and subjected to TCR gene analysis, which revealed oligo or monoclonality for 13 TCRs (**Figure 2d, Extended Data figure 2b**). Five sets of genes encoding TCRαβs directed against the epitopes FLYTYIAKV (FLY-1A), FLYTYIAEV (FLY-1B), EVIGPDGLITV (EVI), MLNYIEQEV (MLN), and AQMWKVVNL (AQM) showed surface-expression by primary T-cells upon retroviral transduction (**Figure 2e, Extended Data figure 2c**). To test whether these TCRs recognize their cognate epitope following endogenous processing and HLA-A2-presentation, a critical go-no go parameter for the respective epitope-TCR pair, we co-cultured TCR T-cells with the TNBC cell line MDA-MB-231 (MM231) that overexpresses either ROPN1 or ROPN1B (**Figure 2f-g**). These tests revealed that FLY-1A and 1B TCRs specifically recognized ROPN1 or ROPN1B^+^ TNBC, respectively (**Figure 2h**). In contrast, MLN- and AQM-specific TCRs did not recognize cancer cells expressing ROPN1/B potentially indicating that these two epitopes are not naturally presented or these TCRs are of a too low affinity for their cognate epitopes (**Figure 2h**). The EVI-specific TCR recognized the parental cancer cells that do not express ROPN1/B (**Figure 2h**) potentially indicating cross-reactivity to other targets. TCRs specific for the EVI, MLN and AQM epitopes were therefore excluded from further characterization.

### TCRs directed against FLY-1A and FLY-1B are highly specific and do not recognize peptides from non-ROPN1/B source proteins

Cross-reactivity of TCR T-cells towards other antigens is a potential safety concern, which when overlooked can result in severe side effects. In example, MAGE-A3 TCR T-cells being cross-reactive to either Titin, a protein highly expressed in cardiomyocytes, or a protein expressed in brain cells, led to patient deaths when tested in clinical trials^33,34^. To assess the extent of cross-reactivity of the FLY-1A and 1B TCRs (**Figure 3a**), we positionally scanned amino acids of the cognate epitopes, replacing each amino acid by any other amino acid to determine which amino acids and which positions are critical for recognition by the corresponding TCR (see **Methods** for details). For both epitopes we found that most amino acid replacements at positions 3-7 resulted in substantial reduction of IFN-γ production by FLY-1A (**Figure 3b**) and FLY-1B (**Figure 3i**) TCR T-cells. With the resulting recognition motif of each TCR (**Figure 3c, j**) an *in silico* search was performed against the complete human proteome to find non-ROPN1 peptides that could theoretically be recognized by either of these two TCRs. Potential peptides coming from this search were shortlisted according to binding to HLA-A2 and recognition by TCR T-cells of peptide-pulsed target cells. These analyses revealed that the FLY-1A TCR recognized 3 non-ROPN1 peptides, namely CLYVFPAKV (CLY), SIWKFPAKL (SIW) and VLFTYVGKA (VLF) coming from the source antigens (E3-independent) E2 Ubiquitin-Conjugating Enzyme (UBE2O), Neuroblastoma Suppressor of Tumorigenicity 1 (NBL1) and Fatty-Acid Amide Hydrolase 1 (FAAH1), respectively (**Figure 3d**). Dose titrations demonstrated that half-maximal responses (i.e., sensitivities) of FLY-1A TCR T-cells for the CLY and SIW peptides are comparable to the cognate FLY-1A epitope, whereas the half-maximal response of this TCR for the VLF peptide is significantly lower compared to the FLY-1A epitope (**Figure 3e**). The amplitudes (i.e., absolute quantities of IFN-γ) for all 3 non-ROPN1 peptides were significantly lower compared to the FLY-1A epitope (**Figure 3e**). We subsequently assessed whether these alternative peptides can be processed and presented by TNBC cells transfected with either one of the three source antigens (**Figure 3g**). Notably, the FLY-1A TCR was unable to recognize TNBC cells overexpressing such source antigens, suggesting that the CLY, SIW and VLF peptides are not a product of the natural antigen processing and presentation machinery (**Figure 3f-g**). The FLY-1B TCR could recognize two non-ROPN1B peptides, namely GMFLYISLA (GMF) and NLYGIVLA (NLY) coming from the source antigens Metal Cation Symporter ZIP8 (SLC39A8) and Sodium/Potassium-Transporting ATPase Subunit Alpha-3 (ATP1A3), respectively (**Figure 3k**). Dose titrations demonstrated that half-maximal responses as well as amplitudes of FLY-1B TCR T-cells for the GMF and NLY peptides were significantly lower or non-existing compared to the FLY-1B epitope (**Figure 3l**). No further investigation of the natural processing and presentation of these peptides was needed.

**Figure 3.**
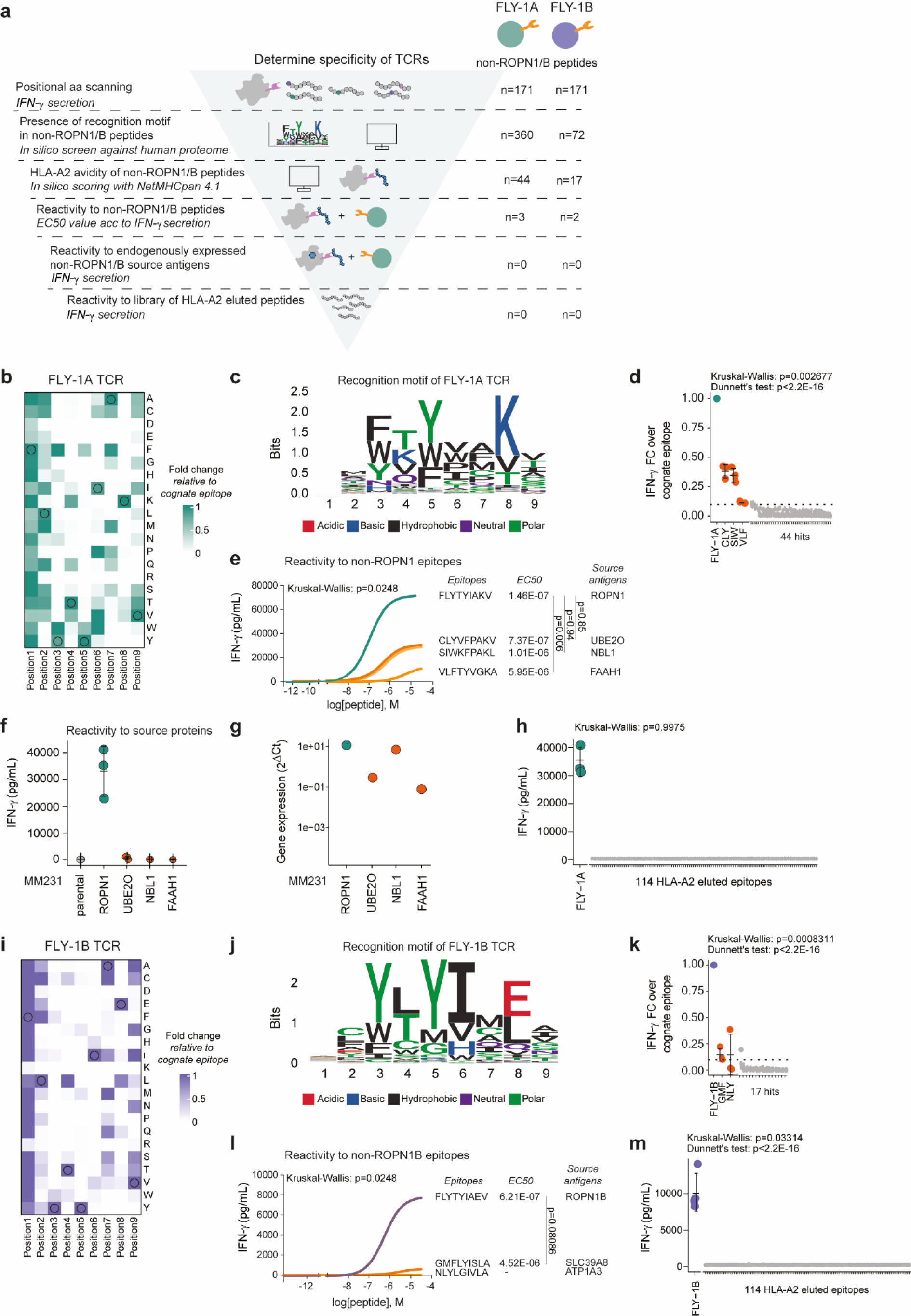
TCRs directed against FLY-1A and FLY-1B have a strict recognition motif and specifically recognize cognate but not alternative epitopes. **a)** Schematic overview of experiments performed to determine TCR specificity (see **Methods** for details). **b)** Heatmap shows relative IFN-γ production by FLY-1A TCR T-cells upon positional amino acid scanning of the cognate epitope. TCR T-cells were co-cultured with BSM cells loaded with single amino acid variants that cover changes in every amino acid at every single position of the cognate epitope (n=171). T-cell IFNγ production is expressed as fold change (FC) compared to the cognate FLY-1A epitope (n=3). Original amino acids from the cognate epitope are circled. **c)** Sequence logo of the recognition motif of FLY-1A TCR T-cells. The height of each letter is scaled in bits using the R package ggseqlogo and represents the probability of that amino acid at that position. The colors of the amino acids represent chemical properties. Motifs were queried against a human protein database using ScanProsite, which yielded 44 potential non-ROPN1 peptides that were predicted to bind to HLA-A2 according to NetMHCpan 4.1. **d)** Dot plot shows IFN-γ production by FLY-1A TCR T-cells upon stimulation with the 44 peptides (10 µg/mL) from **c**, expressed as FC compared to the cognate epitope (n=3). Kruskal-Wallis Rank test was performed followed by Dunnett’s multiple comparisons test: p<2.2E-16. **e)** Representative dose response curves of FLY-1A TCR T-cells that were exposed to non-ROPN1 peptides from **d** with FC>0.1. These peptides included CLYVFPAKV (CLY), SIWKFPAKL (SIW) and VLFTYVGKA (VLF) (depicted in orange); the cognate FLY-1A epitope was included as a comparator (green) (n=3). EC50 values (in M) and source antigens of the three non-ROPN1 peptides are shown on the right side of the plot. Kruskal-Wallis Rank test was performed followed by Dunnett’s multiple comparisons test: CLY vs FLY-1A: p=0.85; SIW vs FLY-1A: p=0.94; VLF vs FLY-1A: p=0.0060. **f)** MM231 cells genetically engineered to express non-ROPN1 source antigens that harbored the peptides from **e** were co-cultured with TCR T-cells to test recognition following endogenous processing and presentation. Dot plot represents absolute IFN-γ levels (in pg/mL) produced by FLY-1A TCR T-cells upon co-culture with MM231 expressing one of the three non-ROPN1 antigens (shown in orange) or ROPN1 (used as a comparator, shown in green) (n=3). **g)** Dot plot represents gene expression of the non-ROPN1 antigens in transfected MM231 cells. Gene expression of the source antigens is depicted as FC relative to GAPDH (2^-ΔCt^) according to qPCR; colors are as in f (n=1). **h)** IFN-γ production by FLY-1A TCR T-cells upon co-culture with BSM cells that were loaded with a library of 114 HLA-A2-eluted peptides. IFNγ production was depicted as FC compared to the cognate epitope (n=3). Kruskal-Wallis: p=0.9975. **i-l)** Specificity testing of FLY-1B TCR T-cells as described in **b-e**. **k**) Dot plot shows IFN-γ production by FLY-1B TCR T-cells upon stimulation with 17 peptides (10 µg/mL) from **j**, expressed as FC compared to the cognate epitope (n=3). Kruskal-Wallis Rank test was performed followed by Dunnett’s multiple comparisons test: p<2.2E-16. **l)** Representative dose response curves of FLY-1B TCR T-cells that were exposed to non-ROPN1B peptides from **l** with FC>0.1. These peptides included GMFLYISLA (GMF) and NLYGIVLA (NLY) (depicted in orange); the cognate FLY-1B epitope was included as a comparator (purple) (n=3). EC50 values (in M) and source antigens of the two non-ROPN1B peptides are shown on the right side of the plot. The EC50 value could not be calculated for the NLY peptide. Data was analyzed using the Wilcoxon signed-rank test: FLY-1B vs GMF: p=0.08086. **m)** IFN-γ production by FLY-1B TCR T-cells upon co-culture with BSM cells that were loaded with a library of 114 HLA-A2-eluted peptides IFNγ production was depicted as FC compared to the cognate epitope (n=3). Kruskal-Wallis Rank test was performed followed by Dunnett’s multiple comparisons test: p<2.2E-16.

Another line of evidence for lack of cross-reactivity came from testing both TCRs for their ability to recognize HLA-A2 eluted non-cognate peptides. To this end, FLY-1A and B TCR T-cells were co-cultured with target cells pulsed with a library of 114 mass spectrometry-observed HLA-A2 eluted peptides, which confirmed non-recognition of any of these peptides for either of the two TCRs (**Figure 3h, m**). Taken together, these findings support the high specificity of both TCRs towards their cognate epitope.

### FLY-1A TCR T-cells effectively kill patient-derived TNBC organoids and outperform standard-of-care treatments

Following our studies into safety, we set out to investigate the killing capacity of ROPN1 specific TCR T-cells using different three-dimensional (3D) TNBC tumoroid models. In a first model we used tumoroids that were derived from the ROPN1 or ROPN1B-overexpressing MM231 cell line (see **Methods** for details). We noted that FLY-1A and FLY-1B TCR T-cells, when loaded on top of these collagen gels, migrated towards and mediated killing of tumoroids expressing ROPN1 and ROPN1B, respectively, over a period of 48h. Notably, FLY-1A TCR T-cells caused 100% tumoroid killing and outperformed cisplatin, which was used as a comparator. FLY-1B TCR T-cells could not completely eradicate the tumoroids within the same time frame and performed similarly to cisplatin (**Figure 4a-b**).

**Figure 4.**
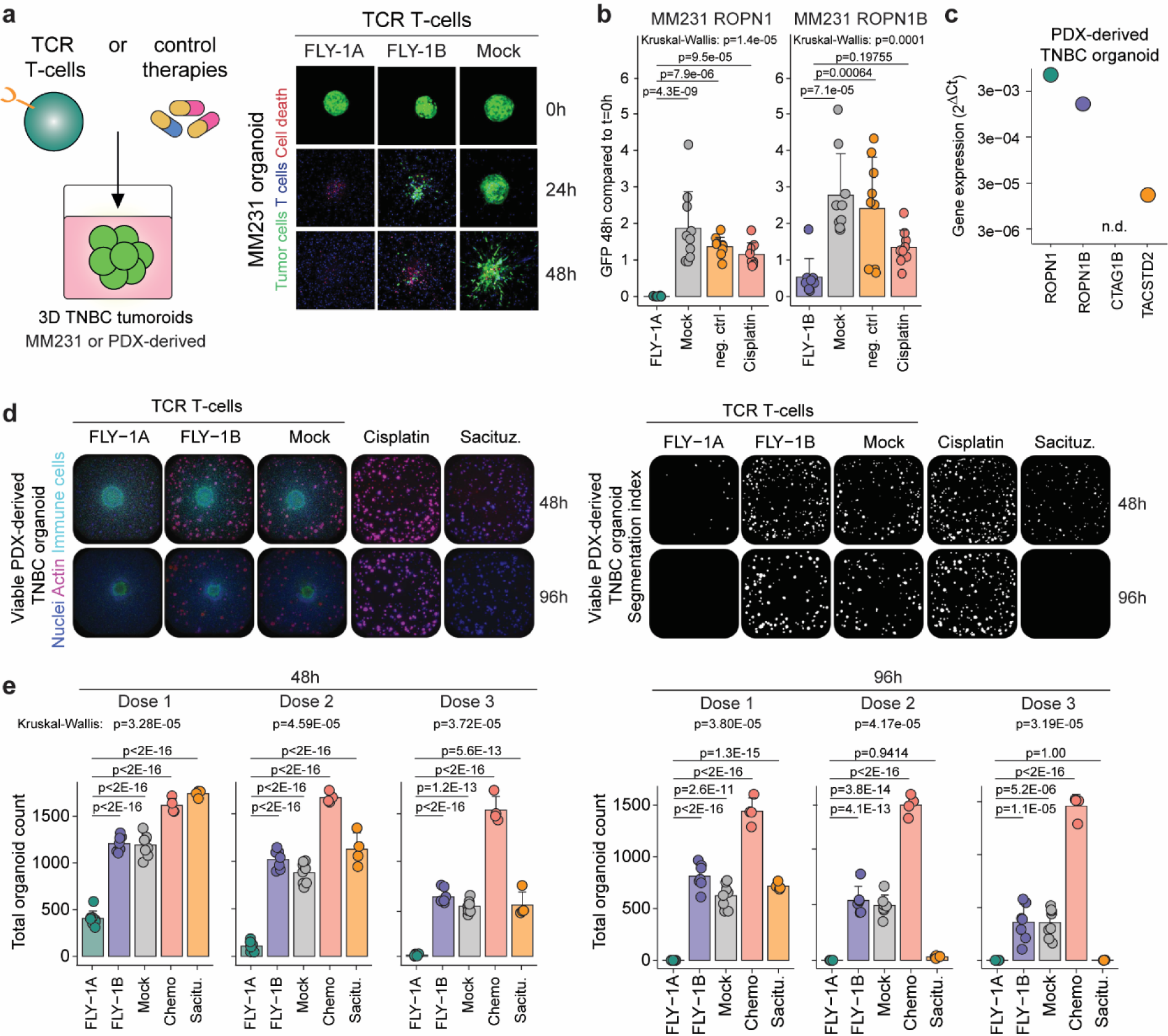
FLY-1A TCR T-cells recognize 3D breast tumoroids and outperform Sacituzumab *in vitro*. **a)** Cartoon depicting the assay set-up to test TCR T-cells or control therapies for their reactivity to three-dimensional (3D) extracellular matrix (ECM)-embedded tumoroids (left panel). Representative confocal fluorescence microscopy images of tumoroids derived from ROPN1/B overexpressing MM231 cells at t=0h, t=24h and t=48h after co-culture with TCR T-cells (n=3 experiments; n=4 replicates per experiment) (right panel). FLY-1A and FLY-1B TCR T-cells were tested against MM231 cells expressing their corresponding antigen, ROPN1 or ROPN1/B. Mock T-cells were included as a negative control. Green color indicates GFP-expressing tumoroid, blue color represents TCR T-cells and red color represents binding by propidium iodide. **b)** Bar graphs display differences in GFP signal from the MM231 ROPN1/B tumoroids 48h after addition of T-cells relative to 0h. Cisplatin or medium (negative control) were used as comparators. **c)** Dot plot represents gene expression of ROPN1 (green), ROPN1B (purple), CTAG1B (NY-ESO1, pink) and TACSTD2 (TROP2, orange) expressed as fold change (FC) relative to GAPDH (2^-ΔCt^) according to qPCR in TNBC tumoroids from PDX (n=1). **d)** Representative images of TNBC PDX-derived tumoroids at 48 and 96h after addition of T-cells. Cisplatin or Sacituzumab govitecan were used as comparators. Left images: Actin shown in pink and nuclei visualized in blue represent living organoids, immune cells are visualized in green. Right images: Segmentation indexes from the images from the left panel representing tumoroid numbers at 48h and 96h. **e)** Bar plots represent total cell count of PDX-derived tumoroid at 48h (left panel) and 96h (right panel) after addition of 3 different doses of the T-cells or drug compounds (n=2 donors, 4 replicates per donor). Abbreviations in figure: PDX, patient-derived xenografts; Sacitu., Sacituzumab govitecan.

In a second series of experiments, we used 3D tumor organoids from TNBC patient-derived xenograft (PDX) tissues that are positive for HLA-A2 and ROPN1 or ROPN1B. These organoids are also positive for TROP2, which is the target for the drug Sacituzumab govitecan, a standard-of-care treatment for TNBC (**Figure 4c**). Again, FLY-1A TCR T-cells were highly efficient in killing the patient-derived organoids that endogenously expressed ROPN1 in a dose-dependent manner and more effectively when compared to the killing of organoids by FLY-1B TCR T-cells after 48h (**Figure 4d-e**). Importantly, FLY-1A TCR T-cells significantly outperformed cisplatin as well as Sacituzumab govitecan. Furthermore, ROPN1-expressing melanoma organoids were also efficiently killed by FLY-1A TCR T-cells to a similar extent as the killing of NY-ESO1-expressing melanoids by NY-ESO1 TCR T-cells (**Extended Data figure 3a-c**).

In line with the superiority of FLY-1A versus FLY-1B TCR T-cells in both organoid models, there is at least a 5-fold difference in effective concentration of cognate epitope that yields half-maximal production of IFNγ (**Figure 3e,l**) pointing to a significantly higher avidity of the FLY-1A TCR T-cells. The TCR specific for FLY-1B was therefore excluded from further characterizations.

### FLY-1A TCR T-cells infiltrate and eradicate TNBC tumors effectively

Next, we assessed tumor cell killing as well as pharmacokinetics of FLY-1A TCR T-cells in mice engrafted human TNBC (see **Methods** and **Figure 5a**). In these experiments, and in line with clinical trials, we preceded the transfer of TCR T-cells with lymphodepleting chemotherapy and supported transferred TCR T-cells with low dose IL-2 (see **Methods** for details). In line with the organoid experiments, treatment with FLY-1A TCR T-cells induced a clear and dose-dependent tumor regression and a single infusion of TCR T-cells significantly outperformed repetitive treatment with Sacituzumab-govitecan (**Figure 5b-c**). The fraction of mice (i.e., 40%) responding to Sacituzumab-govitecan resembled the fraction of patients responding to this drug in clinical trials (i.e., 35%)^3,4^, which together with the abundance of TROP2 in tumor cells used for engraftment (**Figure 5d**), argues that the models we have used are sufficiently sensitive to this antibody-drug-conjugate. In addition, FLY-1A TCR T-cells were detected in dose-dependent quantities in tumors as well as in blood samples (**Figure 5e**). Notably, mice treated with FLY-1A TCR T-cells showed specific enrichment of pMHC^+^ CD8^+^ as well as pMHC^+^ CD4^+^ T-cells within the regressing tumors (**Extended Data Figure 4a-b**). To assess whether the FLY-1A TCR is functionally expressed by CD4^+^ T-cells, we separated FLY-1A TCR CD4^+^ and CD8^+^ T-cells following gene transduction and assessed cytokine production upon co-culture with ROPN1^+^ TNBC cells. We observed that pMHC^+^ CD4^+^ T-cells as well as their pMHC^+^ CD8^+^ counterpart specifically produced IFN-γ and IL-2, suggesting that for its performance the FLY-1A TCR does not require the CD8 co-receptor (**Extended Data Figure 4c**).

**Figure 5.**
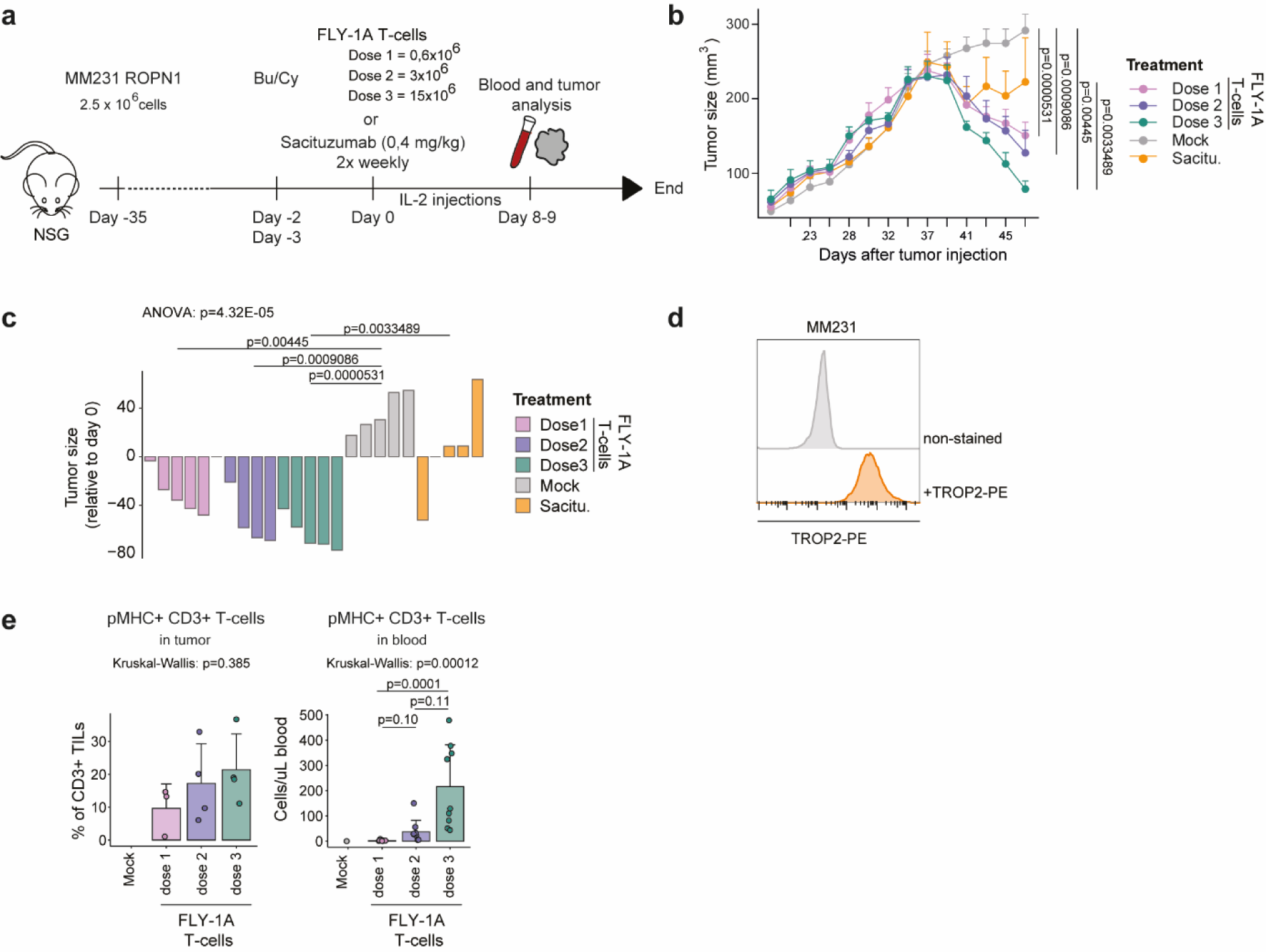
FLY-1A TCR T-cells lead to dose-dependent regression of large TNBC tumors and significantly outperform standard-of-care treatment Sacituzumab govitecan *in vivo*. **a)** Scheme depicting the study design. NSG mice bearing palpable subcutaneous tumors derived from MM231 cells were treated with either 1 transfer of mock or FLY-1A TCR T-cells (0.6, 3 or 15x10^6^ TCR^+^ CD3^+^ T-cells) or Sacituzumab govitecan (0.4 mg/kg) 2x weekly. Blood (n=9 per group) and tumors (n=4 per group) were collected at day 8/9. **b)** Tumor size was measured over time in mice treated with mock T-cells (grey), 3 different doses of FLY-1A TCR T-cells (dose 1: pink; dose 2: purple; dose 3: green) or Sacituzumab govitecan (orange) (n=5 per group). **c)** Waterfall plot representing tumor size at day 11 relative to day -1 per mouse per group (same colors as in b). ANOVA test was performed followed by Tukey post-hoc test. Only significant differences are shown. **d)** Flow cytometric determination of TROP-2 protein expression in MM231 cells stained with TROP2-PE antibody (depicted in orange) or not (negative control, depicted in grey). **e)** Presence of TCR T-cells in tumor (left panel) and blood (right panel). Kruskal-Wallis Rank test was performed followed by Dunn’s multiple comparisons test. Significant differences between mock and other groups was not calculated due to low numbers. Abbreviations in figure: Bu/Cy, busulfan and cyclophosphamide; Sacitu., Sacituzumab govitecan.

### Small scale clinical product of FLY-1A TCR T-cells passes safety and potency test

In a final step, and moving towards clinical development of the FLY-1A TCR, we questioned whether surface expression of this TCR could be enhanced as a function of preferential pairing between TCRα- and β-chains. Upon introduction of exogenous TCRα- and β-chains, there is a possibility of mispairing between transgenic chains and endogenous TCR chains. TCR mispairing could generate new TCR combinations with potentially self-reactive specificities^42,43^, but more importantly (as we have already ruled out recognition of alternative peptides by TCR T-cells, **Figure 3**) it also dilutes the surface expression of transgenic TCRαβ and consequently limits anti-ROPN1 effectivity. To assess the proneness of the FLY-1A TCR to mispairing, we stained FLY-1A TCR T-cells with an antibody specific for its TCR-Vβ13.1 chain as well as with its FLY-1A:HLA-A2 pMHC multimer. We observed equal fractions of TCR T-cells being positive for either TCR-Vβ or pMHC (TCR-Vβ: 31±19% and pMHC: 31±18%, **Figure 6a-b**), suggesting that all surface-expressed FLY-1A TCRβ chains are complexed with all surface-expressed FLY-1A TCRα chains, thereby enabling maximal binding of the cognate epitope coupled to HLA-A2. In fact, the introduction of a cysteine bridge or substitutions of three amino acid residues (LRY), which are reported to reduce mispairing for some TCRs^44,45^, did not result in further improvement of either TCR-Vβ expression or binding to pMHC (**Figure 6c**).

**Figure 6.**
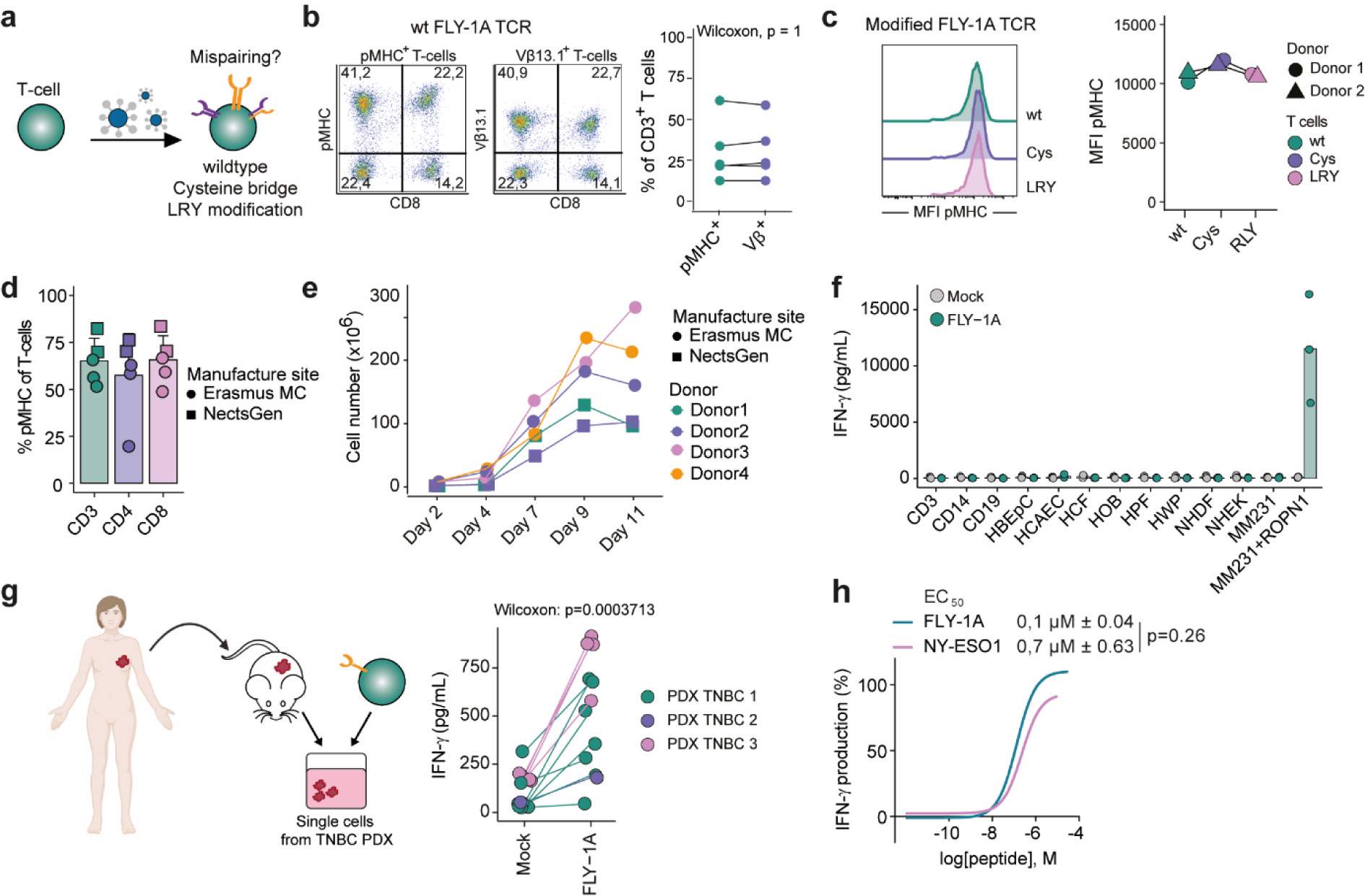
FLY-1A TCR is not prone to mis-pair, and small scale clinical product of non-modified FLY-1A TCR T-cells passes safety and potency assays. **a)** Schematic principle of mis-pairing between transgenic and endogenous TCRα and TCRβ chains following TCR engineering of T-cells. **b)** Representative flow cytometry plot shows FLY-1A wildtype (wt) TCR expression following staining with pMHC (left) or TCRVβ13.1 (middle panel) in CD3^+^ T-cells. The percentages of pMHC (green) and TCR-Vβ (purple) binding T-cells are not statistically significantly different (right panel, ea. n=5 donors). Data was analyzed using the Wilcoxon signed-rank test: p=1. **c)** Representative histogram shows MFI of pMHC^+^ CD3^+^ T-cells transduced with FLY-1A wt TCR (green) compared to FLY-1A TCR with a cysteine bridge (Cys, purplepplure) or LRY modifications (pink) (right panel, n=2 donors). **d)** Bar graph shows FLY-1A wt TCR expression upon staining with pMHC in CD3^+^ (green), CD4^+^ (purple) and CD8^+^ (pink) T-cells following an established GMP process performed in 2 different laboratories (Erasmus MC and NectsGen, ea. n=4 donors). **e)** Cell numbers at manufacturing days 2, 4, 7, 9 and 11 of T-cell products from d. **f)** FLY-A TCR T-cell products were tested for their specificity using a collection of healthy tissue-derived primary cells. IFN-γ production upon stimulation with individual cell types is shown (n=3 donors). ROPN1-expressing MM231 cells were taken along as a positive control. Mock T-cells were used as a negative control. The Wilcoxon signed-rank test per individual healthy cell type: Mock vs FLY-1A: p>0.05. Note: all healthy cell types tested were able to elicit a response by FLY-1A TCR T-cells when pre-loaded with cognate epitope (data not shown). **g)** FLY-A TCR T-cell products from d were tested for their sensitivity using a collection of TNBC PDX. Single TNBC cells were derived from PDX samples that were positive for both ROPN1 and HLA-A2 derived from three different TNBC patients (green, purple and pink represent different TNBC patients). IFN-γ production upon stimulation with single cell suspensions of these TNBC PDX is shown (n=5 T-cell donors). Mock T-cells were used as a negative control. The Wilcoxon signed-rank test: Mock vs FLY-1A: p=0.0003713. **h)** Representative dose response curves of FLY-1A (green, n=6) and NY-ESO1 (pink, n=4) TCR T-cells that were exposed to titrated amounts of their cognate epitope ranging from 1 nM to 30 μM. EC50 values (in μM) are shown above the plot. Data was analyzed using the Wilcoxon signed-rank test: FLY-1A vs NY-ESO1: p=0.26. Abbreviations in figure: Cys, cysteine modification; LRY, LRY modification; MFI, median fluorescent intensity; PDX patient-derived xenografts.

After having determined that the FLY-1A TCR can be considered a so-called ‘dominant TCR’, not needing additional modifications to promote preferential pairing, we manufactured a small-scale FLY-1A TCR T-cell product up to 5x10^8^ cells according to an optimized Good Manufacturing Practice (GMP) process yielding highly effective and juvenile TCR T-cells^46^, currently used in a clinical trial conducted at Erasmus MC (clinicaltrial.gov NCT04729543). These TCR T-cells expanded between 25 to 70-fold in 11 days and contained between 51-82% pMHC^+^ cells within the CD3^+^ T-cell population (**Figure 6d-e**). Importantly, we subjected this clinical-like T-cell product to safety and potency assays that comply with current regulations for first in-human ACT. First, we performed co-culture experiments between the FLY-1A TCR T-cell product and primary cells that were derived from 11 different healthy organs and tissues (**Extended Data table 3**). These experiments demonstrated no elevated IFN-γ levels of the TCR T-cell product when compared to the mock T-cell product (**Figure 6f**), pointing to no or limited off-target toxicity towards healthy tissues. Note that exogenous loading of these primary cells with cognate epitope did evoke a TCR-T-cell IFN-γ response (data not shown). Second, we performed co-culture experiments of the FLY-1A TCR T-cell product with ROPN1^+^/HLA-A2^+^ PDX tissues from three different TNBC patients and one SKCM patient. This FLY-1A TCR T-cell product secreted significantly higher levels of IFN-γ upon co-culture with TNBC PDX (**Figure 6g**) or SKCM PDX single-cell suspensions (**Extended Data figure 3e**) compared to the mock T-cell product, showing that this TCR T-cell product is reactive in an *ex vivo* setting.

## Discussion

We have identified ROPN1, a novel target antigen for ACT with no detectable expression in healthy tissue and high and homogenous expression in over 90% of TNBC, as well as an anti-ROPN1 TCR that demonstrates no cross-reactivity and significant effectiveness in preclinical and near clinical models when compared to standard-of-care drugs. In this study, we have selected the target antigen, epitope as well as the corresponding TCR using a stepwise approach and stringent filtering according to criteria for therapeutic safety as well as efficacy (**Extended Data figure 5**). It is noteworthy that these selections, when applying such criteria, are accompanied by a high number of dropouts. In fact, we started with nearly 300 antigens, 200 epitopes, and >25 TCRs, pointing to a selection stringency, at least for the TCRs, of 10% or less. These numbers may vary for different antigens and different tumor types but do point out that a high number of antigens, epitopes and TCRs are not suited for clinical application and should be identified and dismissed early-on during preclinical development of ACT.

Currently, one of the critical challenges for ACT to treat solid tumors is the lack of suitable tumor-specific targets. Ideally, a suitable target is absent from healthy tissue yet highly abundant in tumor tissue. Starting from a defined set of intracellular antigens, ROPN1 fulfills those criteria, being absent in any of 14 major healthy tissues and highly and homogeneously present in tumor tissues from >300 TNBC patients, even in the metastatic setting and independent of pre-treatments. The only healthy tissues with ROPN1 expression are testis and epididymis, where this protein is expressed in the fibrous sheath of sperm cells^47^, yet these tissues are immune privileged and not present in female TNBC patients. The striking abundance and homogeneity of ROPN1/B in TNBC likely limit the risk of tumor recurrence following ACT since it is reported that outgrowth of antigen-negative tumor cell clones is generally a consequence of low and heterogeneous expression of target antigens within tumors^48–50^. Notably, a recent report highlighted that ROPN1 may act as an oncogene and drive metastasis formation *in vivo* and demonstrated an inverse correlation between ROPN1 expression and survival of TNBC patients^51^, further highlighting the therapeutic value of ROPN1 as a target for ACT.

Moving from target antigen to epitopes, we have applied *in silico* and laboratory tools to select potential T-cell epitopes according to predicted values for peptide-processing and presentation, supplemented with peptides eluted from HLA, and actual binding to HLA-A2. Furthermore, ROPN1 peptides were selected that lack homology to non-ROPN1/B sequences in the human proteome to prevent potential off-target toxicities. Following shortlisting, 11 peptides remained, of which 8 were predicted, 2 were eluted, and 1 peptide (FLY-1A) was both predicted as well as eluted (**Extended Data table 1, 2**). Interestingly, although numbers are limited, there is little concordance between *in silico* predictions and immunopeptidomics. Using a sensitive protocol ^52^, we observed that nine out of the 11 peptides were able to retrieve ROPN1/B-specific T-cell populations from naïve repertoires from healthy donors^42^. From these T-cell populations, we obtained 13 TCRαβs against five epitopes that showed oligo- or monoclonality, of which one TCRαβ against each epitope was surface expressed upon gene transfer into primary human T-cells. When zooming in on T-cell avidity for these five TCRs, we observed a range between 0.1 µM and 4 µM, which is comparable to the avidity observed for the reference TCR directed against NY-ESO1 (**Figure 6h**), demonstrating the feasibility of retrieving highly specific TCRs with natural affinity from naïve repertoires. Next, in a critical go-no go step these TCRs directed against ROPN1/B were challenged for their ability to recognize processed and naturally-presented peptides by TNBC cells. Only the FLY-1A and FLY-1B TCRs survived this test, and the other three TCRs did not, after which the two former TCRs were subjected to extensive safety and efficacy analyses. The FLY-1A and FLY-1B TCRs appeared highly specific for their cognate epitopes according to positional amino acid scanning. Notably, these TCR T-cells failed to recognize non-ROPN1/B peptides that harbored the recognition motif, as was derived from the X-scan, excluding the risk that these TCRs would recognize alternative target antigens. When testing FLY-1A and FLY-1B TCR T-cells for their anti-tumor performance, we demonstrated that FLY-1A TCR T-cells outperformed FLY-1B TCR T-cells with respect to the eradication of ROPN1 or ROPN1B-positive 3D TNBC-as well as melanoma organoids, which is likely linked to their higher functional avidity. In fact, the functional avidity of FLY-1A TCR T-cells is slightly higher compared to NY-ESO1 TCR T-cells (0.1 and 0.7 µM, respectively, **Figure 6h**), with the latter TCR (termed 1G4-α95:LY) already having shown clinical efficacy in treating melanoma and sarcoma patients^53,54^. Furthermore, the FLY-1A TCR is CD8 co-receptor independent, is expressed in both CD4^+^ and CD8^+^ T-cells and FLY-1A TCR-expressing CD4^+^ T-cells produce IFN-γ and IL-2 upon stimulation with epitope (**Extended Data Figure 4c**), characteristics that may aid in engaging CD4^+^ T-cells in an anti-ROPN1 response^55,56^, which are considered critical for inducing durable immune responses. The FLY-1A TCR T-cells showed clear tumor infiltration and effective anti-tumor effects when tested in TNBC-bearing mice. Importantly, the FLY-1A TCR T-cells significantly surpassed the anti-tumor effects seen with standard-of-care drugs, including Cisplatin and Sacituzumab-govitecan, in both tumoroid and mouse models.

Finally, in preparation of a clinical application, we assessed the FLY-1A TCR for its dominance regarding surface expression and demonstrated that the pairing of the transgenic TCRα and β chains could not be improved by cysteine or LRY modifications as described previously^44,57,58^. With the non-modified *wildtype* FLY-1A TCR, we manufactured a clinical-like T-cell product according to an optimized GMP protocol yielding T-cells enriched for a young phenotype (according to CD45RO^-^ and CCR7^+^)^46^, and which protocol is currently used in a clinical trial at Erasmus MC to treat MAGE-C2^+^ melanoma. This FLY-1A TCR T-cell product passed current safety and potency assays as they are used for ACT clinical trials. For example, this TCR T-cell product showed no reactivity towards a range of primary cells derived from healthy organs and tissues, yet showed significant reactivity towards TNBC patient-derived tumor cells.

Even though the preclinical data argue in favor of a safe and effective T-cell product to treat TNBC or melanoma, one cannot exclude that toxicities and/or lack of an anti-tumor response may occur in some patients. With respect to safety, the next step would be to test T-cells gene-engineered with this TCR in a clinical trial according to dose titration. To this end, the preparation of a clinical vector batch and its quality release is being scheduled. With respect to efficacy, we have previously reported that the immunosuppressive microenvironment of TNBC underlies resistance to anti-PD1, and can be grouped according to presence and spatial phenotype of CD8^+^ T-cells ^59^. The excluded phenotype, with CD8^+^ T-cells confined to the tumor margins, harbors dense extracellular matrix and associates with TGFβ pathway activation, whereas the ignored phenotype, with no CD8^+^ T-cells, harbors CD163^+^ myeloid cells and associates with WNT pathway activation. Both these phenotypes, as well as the inflamed phenotype, where CD8^+^ T-cells are present at the tumor center and margins, show similar expressions of ROPN1 (**Extended Data Figure 1d**). Moreover, and in line with our observation that pre-treatment with chemotherapeutic drugs or radiation do not affect ROPN1 expression in TNBC (**Figure 1h**), anti-PD1 treatment did also not affect ROPN1 expression nor did the clinical outcome of this treatment associate with clinical response (**Extended Data Figure 1e**, data from TONIC trial^60^). These preliminary findings suggest that the targeting of ROPN1 may be agnostic for the type of immune suppression as well as the type of pre-treatment.

In summary, we have identified ROPN1 as novel and safe target present in the majority of TNBC patients for ACT and selected the FLY-1A TCR that is highly specific and shows high therapeutic value in advanced tumor models, where it outperforms current treatment options. Currently, T-cells gene-engineered with this TCR are scheduled for their use in a clinical trial to treat patients with TNBC or other ROPN1-positive cancers.

## Methods

### Patient cohorts, databases and code of conduct

TNBC cohort 1: BC with RNAseq (n=347 of which n=66 TNBC, geTMM normalized) accessible through European genome-phenome archive EGAS00001001178 (BASIS cohort)^61^.

TNBC cohort 2: Primary BC with node-negative disease with microarray data (U133) not receiving adjuvant systemic treatment (n=867 of which n=183 TNBC). Data retrieved from gene expression omnibus GSE2034, GSE5327, GSE11121, GSE2990 and GSE7390. Data was normalized using fRMA and batch corrected with ComBat^62^. Details of combined cohorts have been described previously^63^.

TNBC cohort 3: Metastatic TNBC of the Center for Personalized Cancer Treatment (CPCT) cohort with RNAseq (geTMM normalized ^64^) combined and batch corrected with primary TNBC from BASIS ^61^ using the R package ComBat. After batch correction, 22 metastatic TNBC and 66 primary TNBC were used for the analysis.

TNBC cohort 4: Metastatic TNBC from patients treated with anti-PD1 antibody in the TONIC-trial (n=53 of which n=44 paired samples)^60^ with processed transcriptome data of pre- and post-induction treatment biopsies retrieved via controlled access (available through EGAS00001003535).

TCGA: Pan-cancer RNAseq data as well as sample annotation data were retrieved from the USCS Xena browser (n=10,495 of which 1,211 BC, 191 TNBC and 245 SKCM, TPM normalized). The results shown are in part based upon data generated by the TCGA Research Network: https://www.cancer.gov/tcga.

Healthy tissues: RNAseq data of 4 databases covering 66 healthy tissues (Uhlen: n=122 individuals, n=32 tissues^65^; GTEx: n=1,315 individuals, n=53 tissues^66^; Illumina body map: n=32 individuals, n=17 tissues; Snyder Lab: n=25 individuals, n=13 tissues^67^) were downloaded on 13/06/2018 from Expression atlas (TPM normalized).

This study was performed according to the Declaration of Helsinki and the “Code for Proper Secondary Use of Human Tissue in The Netherlands” (version 2002, update 2011) of the Federation of Medical Scientific Societies (FMSF), which aligns with authorized use of coded spare tissue for research. According to national guidelines, no informed consent was required for this study and data as well as ex vivo analysis of spare TNBC tissues was approved by the Medical Ethical Committee of Erasmus MC (MEC.02.953 and MEC-2020-0090, respectively). The collection of patient data and tissues for the generation and distribution of patient organoids was performed in accordance to the guidelines of the European Network of Research Ethics Committees, following European, national and local laws.

### Gene expression: RNAseq, microarray and qPCR

Expression of 276 cancer germline antigens (CGAs, as in Ctdatabase, Ludwig institute, http://www.cta.lncc.br/)^68^ was analyzed in healthy and tumor tissues. Expression of ROPN1 and ROPN1B was evaluated in four different cohorts of healthy tissues and was considered expressed in a tissue when TPM values reached the threshold of >0.2 in at least two cohorts. Expression in tumors (TCGA) was classified as follows: TPM values between 1-9, between 10-100, and >100 were classified as low, moderate, and high expression, respectively. In case of RNAseq data (TNBC cohort 1 and 3) and microarray data (TNBC cohort 2), geTMM and fRMA normalized expression values were used, respectively. We used NY-ESO1 as a reference for expression studies.

Quantitative PCR (qPCR) was performed on cDNA panels of 48 healthy human tissues (OriGene Technologies, Rockville, MD) and TNBC cell lines using MX3000 (Taqman probes ROPN1: Hs00250195_m1; ROPN1B: Hs00250195_m1; CTAG1B (NY-ESO1): Hs00265824_m1; GAPDH: Hs02758991_g1; NBL1: Hs01063631_m1; UBE2O: Hs00222904_m1; FAAH: Hs01038664_m1). Ct values of genes of interest were normalized to GAPDH and relative expression was expressed as 2^-ΔCt^.

### Protein expression: immunohistochemical staining

Stainings were performed using large cores of healthy tissues (2 mm in diameter) covering 16 major tissues ea. from 2-6 individuals (derived from autopsy or resection, n=62) as well as FFPE tissue microarrays of TNBC primary tumor tissue covering 311 patients^69^. In addition, whole tissue sections of paired primary and metastatic TNBC tumors (n=15) were stained^59^. Staining with anti-ROPN1 antibody (which detects both ROPN1 and ROPN1B protein, 1:200, HPA052530, Atlas) or NY-ESO1 antibody was performed following heat-induced antigen retrieval for 20 min at 95°C. After cooling to RT, staining was visualized by the anti-mouse EnVision+® System-HRP (DAB) (DakoCytomation, Glostrup, Denmark). Human testis tissue was used as positive control tissue. Stainings were manually scored on intensity and percentage of positive tumor cells using Distiller (SlidePath) software independently by 3 investigators (MT, DH, DK).

### Generation of ROPN1/B expressing TNBC cell line

To generate a ROPN1/B-overexpressing triple negative breast cancer (TNBC) cell line, a ROPN1 or ROPN1B-GFP fragment (amino acid sequence accessible under UniProtKB Acc. No. Q9HAT0/Q9BZX4) (ROPN1/B-2A-GFP) was ordered via GeneArt (Regensburg, Germany) and cloned into the PiggyBac vector (a kind gift from Dr. P.J. French, Erasmus MC, Rotterdam, the Netherlands) using In Fusion cloning kit (Takara). Subsequently, the MDA-MB-231 cell line (MM231, ECACC catalogue no. 92020424, a cell line model for TNBC) was stably transfected with PiggyBac ROPN1/B-GFP DNA using Lipofectamine P3000 transfection reagent kit (Invitrogen) and Transposase Expression vector DNA (System Biosciences). Cells were cultured with 2 µg/mL puromycin (Life technologies). Expression of ROPN1/B and GFP was confirmed with qPCR and flow cytometry, respectively.

### Identification, selection and ranking of ROPN1/B peptides

ROPN1/B peptides were ranked according to multiple in silico methods to predict different aspects of immune reactivity^70^ (i.e., NetMHCpan^71^; NetCTLpan^72^; SYFPEITHI^73^; and RANKPEP^74–76^). For details on predicted features of ea. Of these algorithms see Hammerl et al.^70^. For immunopeptidomics, the ROPN1/B-expressing MM231 cells (3x10^8^) were treated with 50 pg/mL human recombinant IFN-γ (Preprotech) for 24h and harvested using EDTA before immunoprecipitation of MHC class I molecules. Peptides were eluted and measured with mass spectrometry as before. In addition, 1x10^8^ ROPN1B-expressing K562ABC cells were treated with epigenetic drugs (treatment regimen as described in^77^) prior to immunoprecipitation. K562ABC cells are lymphoblastic cells that express HLA-A, B and C alleles and are typically used for antigen presentation research; the ROPN1B-expressing variant of K562ABC cells was generated as described for ROPN1/B-expressing MM231 cells. Peptides were eluted and measured with mass spectrometry as described previously^39,78^ (also see **Supplementary methods**). Lastly, peptides were searched in available databases of peptides eluted from tumor and healthy tissues as well as cell lines (https://www.zhang-lab.org/caatlas/). The top-10 predicted peptides per in silico tool as well as unique peptides (not overlapping with the predicted peptides) retrieved from immunopeptidomics were checked for cross-reactivity with Expitope^41^. Peptides that overlapped with peptides from non-ROPN1/B human proteins with up to 2 amino acid mismatches (i.e., 7 out of 9 amino acids are identical) were excluded from further analysis. The final list of peptides was ordered at ThinkPeptides (ProImmune, Oxford, United Kingdom), dissolved in 50-75% DMSO and stored at -20°C until use. The HLA-A2 stabilization assay was performed using T2 cells as described in^79^ with a slightly adapted protocol (see **Supplementary methods**). In brief, T2 cells were exposed to titrated amounts of peptide (range from: 31 nM to 31 μM, n=3) and HLA-A2 expression using an HLA-A2 PE antibody (clone BB7.2, 1:20, Abcam) was measured using flow cytometry (FACS Celesta, BD) and analyzed using FlowJo software (version 10.7.1, TreeStar, Ashland, Oregon, USA).

We calculated two parameters of binding avidity to HLA-A2: (1) amplitude, which was the difference in fluorescence intensity between the highest concentration and baseline; and (2) half-maximal effective concentration (EC50), which is the calculated concentration of peptide (using GraphPad Prism 5.0) giving half-maximal response. Eleven peptides adhered to the following 2 criteria (1) EC50 of <1x10^-4^ M; and (2) binding amplitude of >0.5 relative to the reference peptide gp100^YLE^, and these peptides were ranked according to amplitude values.

### Enrichment of ROPN1/B epitope-specific CD8^+^ T-cells

Enrichment of epitope-specific T-cells was performed by co-culturing naïve T-cells with autologous CD11c^+^ cells that were loaded with the shortlisted peptides using PBMCs from 2 to 7 healthy donors (see **Supplementary methods**) per tested peptide. Following the first co-culture cycle, ROPN1-specific T-cells were amplified over another three cycles using peptide-pulsed autologous PBMC supplemented with medium containing common-γ cytokines according to a protocol we have previously^52^. Following enrichment, T-cells were tested for ROPN1 and/or ROPN1B epitope-specific IFN-γ production. To this end, supernatant from overnight co-cultures of T-cells with T2 cells (1x10^6^/mL) that were pulsed with epitope (10 ng/mL) was collected and IFN-γ production was measured with an enzyme-linked immunosorbent assay (ELISA, BioLegend) according to the manufacturer’s protocol. T2 cells loaded with an irrelevant epitope were included as a negative control. T-cell IFN-γ production was considered epitope-specific in case levels exceeded 200 pg/mL, and levels were minimally twice as high as for irrelevant epitope. T-cells that fulfilled these criteria were stained with peptide:MHC (pMHC) tetramers (Tetramer shop, Kongens Lyngby, Denmark) to determine the frequency of epitope-specific T-cells. Events were acquired with FACS Celesta and analyzed using FlowJo software. In case T-cell binding of pMHC was observed in more than 0.5% of CD3^+^ T-cells, these cells were FACS-sorted with pMHC multimers.

### TCR cloning and sequence identification

CD8^+^ T-cells were exposed to the SMARTerTM RACE cDNA Amplification Kit (Clontech/Takara) to identify and annotate ROPN1/B epitope-specific TCRα- and β-chains based on Kunert et al. ^77^. Using the IMGT database and the HighV-QUEST tool (http://www.imgt.org), the TCR-V, D, and J sequences were annotated according to the Lefranc nomenclature^80,81^. In case for a given T-cell population, TCRα sequences represented 30% or more of all functional sequences of TCRα and the same for TCRβ (i.e., >30% clonal sequences of both TCRα and TCRβ), then those TCR chains were matched with each other. These sets of TCRα and TCRβs were codon optimized (GeneArt, Regensburg, Germany) and cloned into the pMP71 vector (a kind gift of prof. Wolfgang Uckert, MDC, Berlin, Germany) using a TCRβ-2A-TCRα cassette.

### TCR gene transfer

Upon activation with anti-CD3 Monoclonal Antibody (OKT3, Thermofisher), PBMCs from healthy donors were transduced with TCR-encoding retroviruses (pMP71) or empty vector that were produced by a co-culture of 293T and Phoenix-Ampho packaging cells, as described previously ^82,83^. Staining for surface-expressed TCR transgenes was performed using pMHC dextramers (Immudex) as described in ^84^. In case pMHC complexes were insensitive to detect TCR T-cells, we used antibodies directed against TCR-Vβ (Beckman Coulter) and CD137-APC (1:25, BD). The expression of CD137 was measured following 48h stimulation with epitope-loaded BSM melanoma cells. In case pMHC or CD137 expression in more than 5% of CD3^+^ T-cells was observed in at least 2 donors, TCR T-cells were then MACS-sorted using pMHC complexes or FACS-sorted according to upregulated CD137 expression.

### Co-culture experiments of TCR T-cells with target cells

TCR-transduced T-cells (6×10^4^/well in a 96-well round bottom tissue culture treated (TCT) plate) were co-cultured with target cells (2×10^4^/well) in a total volume of 200 μL RPMI medium supplemented with 25 mM HEPES, 6% human serum (Sanquin, Amsterdam, the Netherlands), 200 mM L-glutamine, and 1% antibiotics for 24h at 37°C/5%CO_2_. Parental or ROPN1/B expressing MM231 tumor cells were pre-treated 48h with IFN-γ prior to co-culture. T-cell recognition of endogenously processed and presented epitopes was demonstrated in case IFN-γ levels exceeded 200 pg/mL, and levels were minimally twice as high as for mock T-cells. To test sensitivity of FLY-1A and FLY-1B TCR T-cells, T-cells were co-cultured with BSM cells loaded with epitope concentrations ranging from 1 pM to 30 μM to determine EC50 values.

### Positional amino acid scanning of ROPN1/B epitopes

Positional amino acid scanning of ROPN1/B epitopes was performed to assess the recognition motifs of TCRs. TCR T-cells were co-cultured with BSM cells that were pulsed with alternative peptides (i.e., 10 µg/mL) covering all individual amino acids as replacements at every single position in the cognate ROPN1/B epitope (n=171). Critical amino acid substitutions were defined as those that resulted in at least a 90% drop in IFN-γ production when compared to the cognate epitope. Once having identified alternative peptides that contained such critical amino acid(s), we could assign a fold change value (FC) for every amino acid at every position relative to the cognate epitope. We calculated the frequency per amino acid at every position by dividing the FC of each amino acid by the sum of FCs of all amino acids per position.

The resulting frequency matrix was used as input to create the sequence logo presented in bits with the R package ggseqlogo^85^. The resulting recognition motif was scanned for its occurrence in the human proteome using the ScanProsite tool^86^. Non-ROPN1/B peptides that contained the recognition motif were scored for their *in silico* binding to HLA-A2 with the NetMHCpan 4.1 software^87^. HLA-A2 binders (weak as well as strong binders) were subsequently synthesized and tested for their capability to induce responses by ROPN1/B TCR T-cells. To this end, alternative peptides were loaded onto BSM cells and tested at a single dose of 10 µg/mL, and those that resulted in T-cell IFN-γ secretion (FC>0.1 compared to the cognate epitope) were used to determine EC50 values according to dose titrations. Importantly, non-ROPN1/B source antigens that harbor such peptides were overexpressed in MM231 cells, after which these MM231 cells were co-cultured with TCR T-cells. Only in case there was no reactivity by TCR T-cells towards alternative source antigens, the peptide was considered not to be a product of endogenous processing and presentation. Finally, TCR T-cells were screened for lack of reactivity towards 114 HLA-A2-eluted non-cognate peptides as described in^88^.

### Testing reactivity of TCR T-cells towards 3D organoid models

TCR T-cells were subjected to tracking and monitoring in three-dimensional (3D) tumoroid models of ROPN1/B-GFP expressing MM231 cells, and patient-derived organoids (PDX) from breast and skin tumors. To this end, we used image-guided injection of droplets with cells into a collagen gel to form extracellular matrix (ECM)-embedded tumoroids as described previously ^89–91^. Overnight, gels were overlayed with T-cell medium containing 0.4 µM propidium iodide (PI). Subsequently, TCR or mock T-cells were labeled with 0.1 µg/mL Hoechst 33242 (Invitrogen) at 37°C/5%CO_2_ for 1h and 50,000 labelled T-cells were dispersed on top of the collagen gel. Some tumoroids were treated with 20 µM cisplatin as a positive control for tumor cell killing. Tumoroids were imaged at indicated time points by confocal microscopy using a Nikon TE2000 equipped with a Prior stage controlled by NIS Element Software, and a temperature and CO_2_-controlled incubator. 10 µm Z-stacks were captured spanning the entire tumoroid using a 20x objective. Images were stitched where necessary and analyzed by CellProfiler (v 2.2.0). A maximum projection was generated summing Hoechst, GFP, and PI signals in each image.

Organoid models from PDX from breast and skin tumor-derived were established as described ^92^. PDX organoids (BR5010B & ME12086B) were seeded in 3D hydrogel (Proprietary, Crown Bioscience Netherlands B.V.) in 384-well plates (Greiner Bio-One B.V.) and incubated for 48h. TCR T-cells were stained with cell tracker (CellTracker™ Green CMFDA Dye, Invitrogen), resuspended in T-cell medium and added on top of the 3D hydrogel at three different ratios of organoids to T-cells. Cisplatin and Sacituzumab-govitecan were added to tumor organoids at 0.1, 1 and 10 µM and 0.1, 1 and 10 nM, respectively. After 48 and 96h, plates were fixed, stained and tumor volumes were analyzed. Imaging was performed using Molecular Devices ImageXpress Micro XLS (Molecular Devices) with a 4x NIKON objective. Image analysis was performed using Ominer® software, and experiments and analyses were performed by personnel at the premises of Crown Bioscience, Netherlands, Leiden (see **Supplementary methods**).

### In vivo performance of TCR T-cells in a TNBC xenograft model

ROPN1 expressing MM231 tumor cells (2,5x10^6^) were suspended in matrigel and subcutaneously (sc) transplanted in the right flank of NSG mice (NOD.Cg-Prkdcˢᶜⁱᵈ Il2rgᵗᵐ¹ᵂʲˡ/SzJ, Charles River Laboratories, Paris, France). Tumor engraftment and treatment regimen was carried out as described previously^93^ with only minor changes. In short, mice (n=9 per treatment group) received a single intravenous (iv) transfer of 0.6, 3 or 15×10^6^ human TCR or mock T-cells followed by sc IL-2 injections (1x10^5^ IU) for eight consecutive days following T-cell transfer. Alternatively, mice received two iv injections per week of Sacituzumab govitecan (Trodelvy, Gilead Sciences Ireland UC; 0.4 mg/kg according to the manufacturer’s instructions). At day 11, tumor regressions were measured relative to day 0, and tumor regression in mice treated with TCR T-cells was compared with mock T-cells and Sacituzumab govitecan. In addition, blood (n=9 mice) and tumor (n=4 mice) samples were collected at day 8/9 to detect transferred T-cells with flow cytometry. Single cell suspensions were obtained from the tumors using collagenase treatment as described above. Cells (either from blood or tumor) were washed with PBS and incubated with pMHC and additional antibodies (CD3-BV421, 1:20, BD; CD8-BV650, 1:100, BD; CD4-V500, 1:50, BD; CD137-APC, 1:25, BD; CD279-APC-Cy7, 1:50, ITK; and CD366-PE-Cy7, 1:10, BD). Events were acquired with FACS Celesta and analyzed using FlowJo and R packages ggplot2 and ComplexHeatmap^94,95^.

### Assessing preferential pairing of TCR chains

The mispairing between the transgenic and endogenous TCR genes was assessed by transducing T-cells with the wildtype or modified TCRαβ chains or the TCRβ chain only. We applied two separate modifications of TCR chains, namely the introduction of a cysteine bridge or the LRY amino acids, which have both been reported to enhance preferential pairing between the transgenic TCRα and TCRβ chains^44,45^. TCR expression was assessed by Flow Cytometry upon staining with pMHC multimers and Vβ13.1 antibody.

### GMP manufacturing of T-cell product

We made the TCR T-cell product using a Good Manufacturing Practice protocol that has been optimized previously^4640^ and implemented in a clinical trial (clinicaltrials.gov identifier: NCT04729543). To this end, frozen PBMCs were thawn and resuspended in T-cell medium to a concentration of a 1x10^6^ cells/mL. Cells were activated with soluble anti-CD3 (30 ng/mL, ThermoFisher) and anti-CD28 (30 ng/mL, Life technologies), IL-15 (110 IU/mL, Miltenyi Biotec) and IL-21 (0.1 IU/mL, Miltenyi Biotec) for 48h. Retronectin-coated (overnight, 12 µg/mL, Takara) Non Tissue Culture flat bottom plates (24 well) were blocked with 1 mL PBS/2%FBS, after which 0.3 mL retroviral supernatant supplemented with IL-15/IL-21 was added per well. Plates were centrifuged for 15 min at 1,000g (slow brake). T-cells (1x10^6^/well) were resuspended in 0.3 mL retroviral supernatant and added to each well. Plates were centrifuged for 60 min at 1,000g (slow brake) and incubated at 37°C/5%CO_2_. After 5h, 800 µl/well T-cell medium supplemented with IL-15/IL-21 was added to each well and incubated overnight, after which 1.2mL supernatant was removed and 0.6mL fresh retroviral supernatant supplemented with IL-15/IL-21 was added per well. Again, plates were centrifuged to prepare for a second hit with retroviral supernatant, as described above. T-cells were then harvested, and cultured at a concentration of 0.25x10^6^/mL in T-cell medium supplemented with IL-15/IL-21 for 3-6 days. T-cells were either directly used in co-culture assays or frozen down. Manufacturing was performed at Erasmus MC, Rotterdam and at the NectsGen facility, Leiden.

### Reactivity of TCR T-cells towards healthy cells

Primary human healthy cells from HLA-A2^+^ donors were obtained via ScienCell or Promocell (**Extended Data table 3**) and cultured and frozen down according to the manufacturer’s instructions. On the day of assay, primary cells and TCR T-cells were thawn and rested for 1 and 2h, respectively, after which co-culture assays were performed as described above.

### Reactivity of TCR T-cells towards patient-derived xenografts in vitro

We assessed T-cell reactivity towards TNBC and melanoma-derived PDX. PDX’s were maintained using serial transplantation using NSG mice. Tumors, once isolated, were cut into smaller fragments in DMEM complete medium and washed with PBS. Cells were resuspended in fresh Collagenase A solution (Sigma) in PBS (1 mg/mL) and incubated for 45 min at 37ᵒC/5%CO_2_ while being regularly mixed. EDTA (0.1mM) was added and cells were pushed through a 70 µM cell strainer. Single cells were either pre-treated with IFN-γ for 24h or used directly in co-culture assays. Mock or TCR T-cells (1x10^5^) were co-cultured with single tumor cells in a 1:1 ratio overnight after which supernatant was harvested to detect IFN-γ.

### Statistical analyses

Statistical analysis was performed with GraphPad Prism 5.0 and Rstudio. Statistically significant differences between 2 groups were calculated with Mann-Whitney U test for independent samples or the Wilcoxon signed-rank test for paired samples. To determine statistically significant differences between multiple groups, one-way analysis of variance (ANOVA) with adjustments for multiple comparisons with Tukey’s post-hoc test was performed. If assumptions for ANOVA were not met, the Kruskal-Wallis Rank test was performed followed by Dunnett’s or Dunn’s multiple comparisons test. Differences were considered statistically significant when p < 0.05.

## Acknowledgment

We thank the FACS Sorting Shared Facility (FSSF) of Erasmus MC, especially Andrea Sacchetti, Harm de Wit and Peter van Geelen, for their assistance. This work was supported by the Health Holland Public-Private-Partnership award EMC-TKI LSH20020, the Erasmus MC Daniel den Hoed Foundation and the Dutch Cancer Society (KWF; project no. 2014-7087).

## Author contributions

DH and RD conceived and supervised the project. DK, DH and RD conceptualized and designed experiments, further supported by JWM and RJMA. DK, DH, MvB, RW, DR, KK performed T-cell based experiments and analyzed data. MT, AT, RF performed immune stainings. CYL and EHJD performed organoid experiments and analyzed the data. JM, MdB, SIB, JAD, MBS generated and analyzed MS data. JWM and MK provided translational data of patient cohorts. DK, DH and RD wrote the manuscript. All authors read and reviewed the manuscript.

## Competing interests

This research was conducted according to the requirements of objectivity and integrity standards. DH is listed as inventor for European patent application no. P128827EP00. MK has received research support from BMS, Roche, AstraZeneca, personal fees from AstraZeneca, Daiichi Sankyo, Domain Therapeutics, Alderaan, BMS, MSD, Gilead, Roche outside the submitted work (all paid to the NKI/AVL). RD has received research support from MSD and Bayer, personal fees from Bluebird Bio, Genticel, other support from Pan Cancer T B.V. outside the submitted work (all paid to the Erasmus MC Cancer Institute) and is listed as inventor for European patent application no’s. P130556EP00 and P128827EP00. All other authors (DK, MvB, RW, DR, KK, MT, CYL, ATJ, RF, JM, MdB, SIB, JAD, MK, EHJD, MBS, JWM, RJMA) declare no competing interests.

## Extended Data

**Extended Data figure 1.**
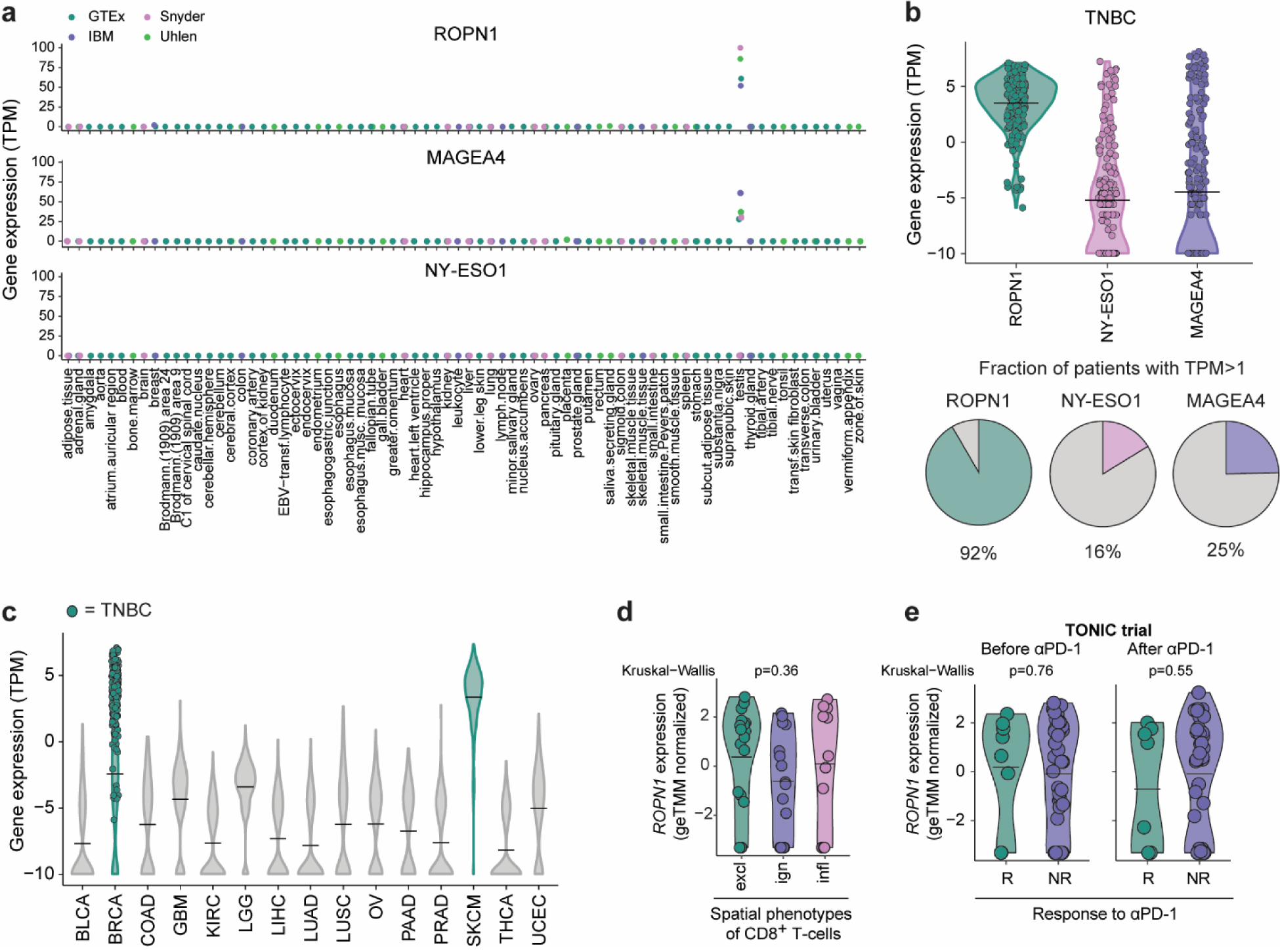
Expressions of ROPN1/B, NY-ESO1 and MAGE-A4 target antigens in healthy tissues, TNBC and various tumor types. **a)** Dot plots represent average gene expression of ROPN1, MAGEA4 and CTAG1B (NY-ESO1) in healthy tissues (TPM values) according to RNAseq of 4 different healthy tissue datasets (Uhlen: n=122 individuals, n=32 tissues (purple); GTEx: n=1,315 individuals, n=53 tissues (dark green); Illumina body map: n=32 individuals, n=17 tissues (light green); Snyder Lab: n=25 individuals, n=13 tissues (pink)). **b)** Violin plot shows distribution of gene expressions of ROPN1 (green), MAGEA4 (purple) and CTAG1B (NY-ESO1, pink) in the TCGA TNBC cohort (n=191, TPM normalized). Pie charts show fractions of TNBC tumors with gene expression for these 3 target antigens (TPM>1). **c)** Violin plot shows distribution of ROPN1 gene expressions in 15 tumor types retrieved from TCGA cohorts (TPM normalized). BRCA and SKCM are visualized in green; individual TNBC samples are shown as green dots. **d)** Violin plot shows distribution of ROPN1 gene expression according to spatial phenotypes of CD8^+^ T-cells of metastatic TNBC retrieved from the TONIC cohort (n=53, geTMM normalized). The spatial phenotypes of CD8^+^ T-cells included: excluded (excl), ignored (ign) and inflamed (infl) as described in ^59^. Kruskal-Wallis Rank test was performed between spatial phenotypes: p=0.36. **e)** Violin plot shows distribution of ROPN1 gene expression in responders (R) versus non-responders (NR) before and after αPD-1 treatment in metastatic TNBC retrieved from the TONIC cohort (n=53, geTMM normalized). Kruskal-Wallis Rank test was performed between R and NR: before αPD-1: p=0.76, after αPD-1: p=0.55. Abbreviations in figure: BLCA, Bladder Urothelial Carcinoma; BRCA, Breast Carcinoma; COAD, Colon Adenocarcinoma; excl, excluded; GBM, Glioblastoma Multiforme; ign, ignored; infl, inflamed; KIRC, Kidney Renal Clear Cell Carcinoma; LIHC, Liver Hepatocellular Carcinoma; LUAD, Lung Adenocarcinoma; LUSC, Lung Squamous Cell Carcinoma; OV, Ovarian Serous Cystadenocarcinoma; PAAD, Pancreatic Adenocarcinoma; PRAD, Prostate Adenocarcinoma; SKCM, Skin Cutaneous Melanoma; THCA, Thyroid Carcinoma; UCEC, Uterine Corpus Endometrial Carcinoma.

**Extended Data figure 2.**
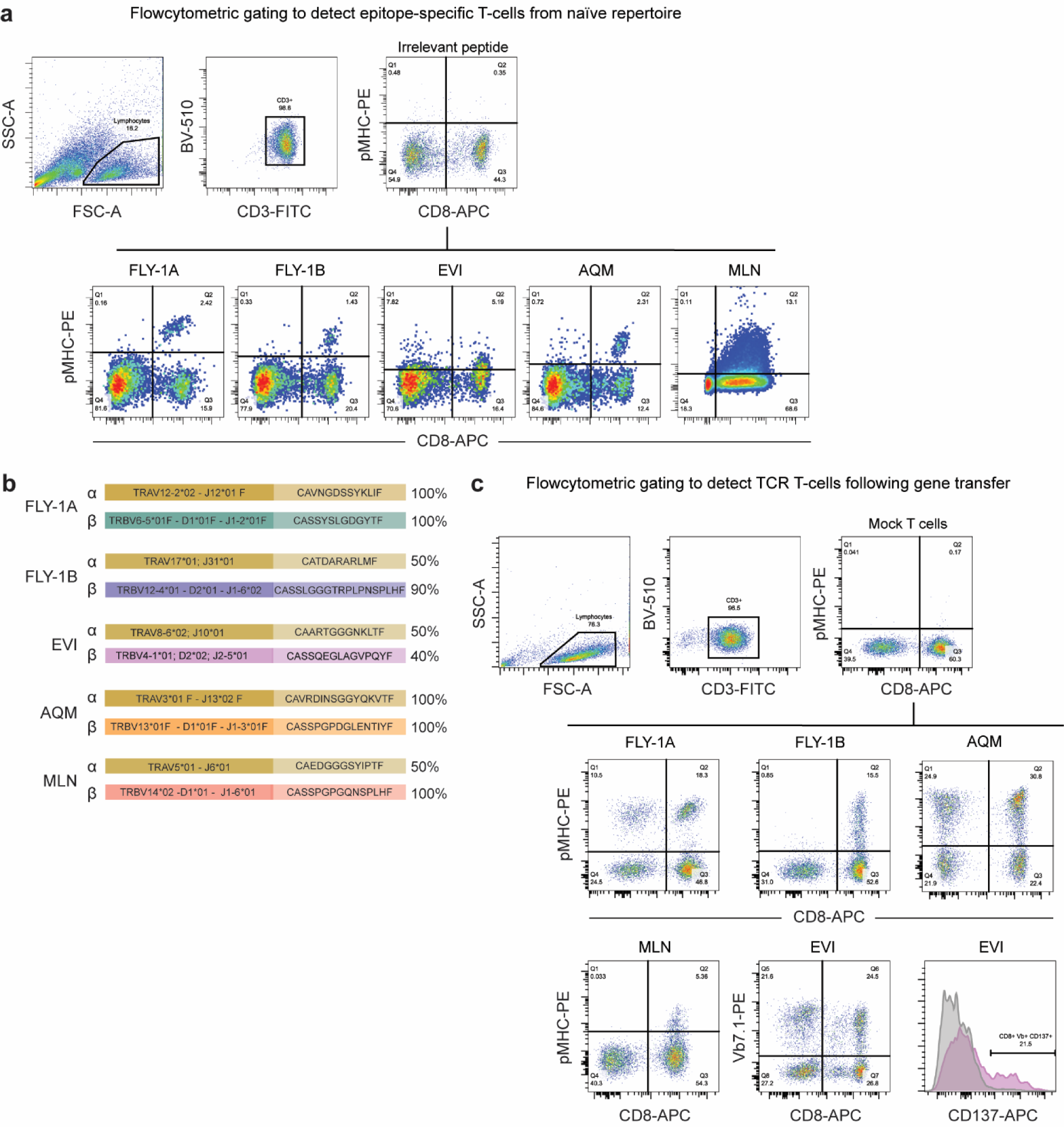
Retrieval of ROPN1/B-specific T-cells and TCR genes. **a)** Illustration of strategy employed for flow cytometry of pMHC binding of T-cells following 4 cycles of co-culture with epitope-loaded APCs (i.e., T-cell enrichment for epitope-specific T-cells). Upper plots show gating strategy to define epitope-specific T-cells according to pMHC staining of T-cells that were enriched using an irrelevant epitope. Lower five plots are representative pMHC stainings of T-cells that were enriched with FLY-1A, FLY-1B, EVI, AQM or MLN epitope, respectively. **b)** Identification of T-cell receptor V-alpha (TRAV and J according to IMGT nomenclature), T-cell receptor V-beta genes (TRBV, D and J; blue) and T-cell receptor C genes (with starting and ending amino acids) cloned from T-cells from a; percentage reflects fraction of these sequences among all identified TCR sequences per epitope. **c)** Representative flow cytometry plot of pMHC binding of TCRs from b following gene transfer into T-cells (ea. n=2 donors). Note: in case of EVI TCR, the specific pMHC complexes appeared insensitive in detecting TCR T-cells, and were replaced by stainings with antibodies directed against TCR-Vβ7.1 and CD137 (the latter following 48h stimulation with cognate epitope-loaded BSM cells).

**Extended Data figure 3.**
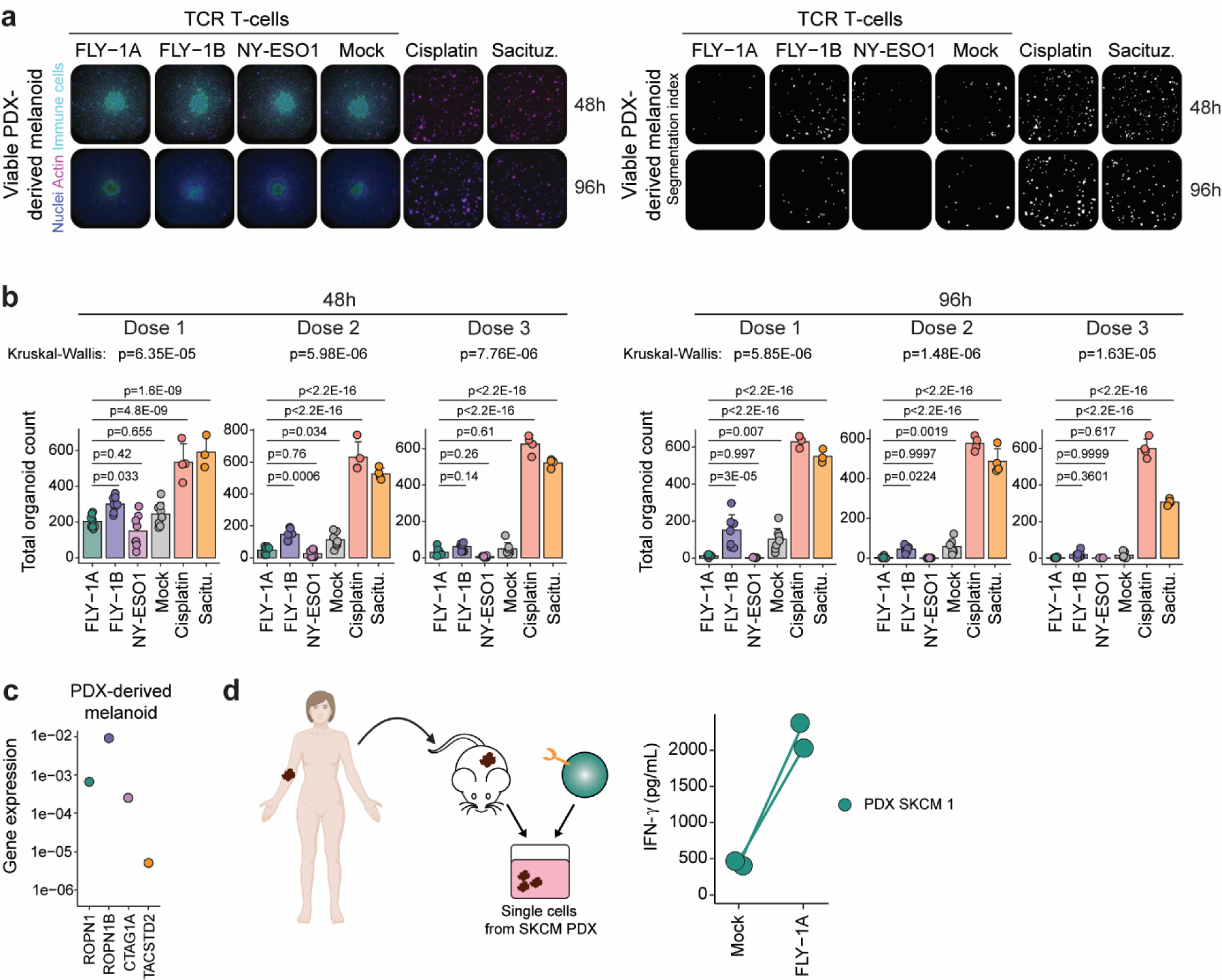
FLY-1A TCR T-cells recognize melanoma tumoroids and patient-derived melanoma cells. **a)** Representative microscopy images of melanoma PDX derived tumoroids at 48h and 96h after addition of T-cells or drug compounds (See **Methods** and legend to Figure 4 for details). Left images: Actin shown in pink and nuclei visualized in blue represent living melanoma organoids, immune cells are visualized in green. Right images: Segmentation indexes from the images from the left panel representing tumor cell numbers at 48 and 96h. **b)** Bar plots represent total cell count of PDX-derived tumoroid at 48 (left panel) and 96h (right panel) after addition of 3 doses of the T-cells or drug compounds (n=2 donors, 4 replicates per donor). **c)** Dot plot represents gene expressions of ROPN1 (green), ROPN1B (purple), CTAG1B (NY-ESO1, pink) and TACSTD2 (encoding for TROP2, orange) of a melanoma PDX-derived tumoroid expressed as fold change relative to GAPDH (2^-ΔCt^) according to qPCR (n=1). **d)** FLY-1A TCR T-cells were tested for their sensitivity using SKCM PDX. Single melanoma cells were derived from 1 PDX sample that was positive for both ROPN1 and HLA-A2. IFN-γ production upon stimulation with single cell suspensions of this PDX is shown (n=2 donors). Mock T-cells were used as a negative control. Abbreviations in figure: PDX, patient-derived xenografts; Sacituz, Sacituzumab govitecan; SKCM, Skin Cutaneous Melanoma.

**Extended Data figure 4.**
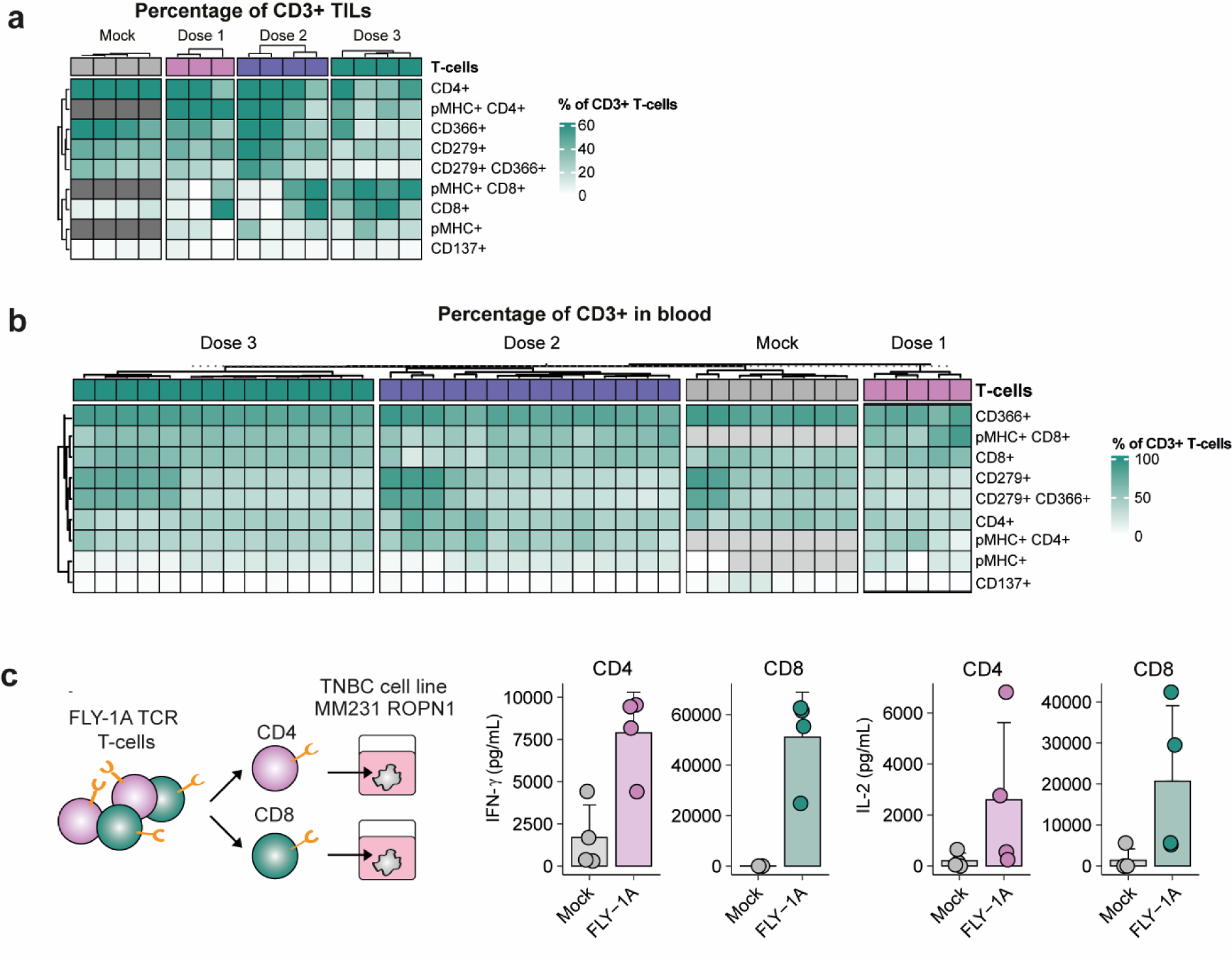
FLY1A^+^ CD4^+^ T-cells recognize ROPN1 TNBC. **a)** Flow cytometric analysis of T-cells retrieved from tumors at day 8/9 following ACT into tumor-bearing mice (See **Methods** and legend to Figure 5 for details). Heatmap reflects percentages of the following markers: pMHC^+^, CD4^+^, CD8^+^, pMHC^+^CD4^+^, pMHC^+^CD8^+^, CD137^+^, CD279^+^, CD366^+^ and CD279^+^CD366^+^ within CD3^+^ T-cells. **b)** Flow cytometric analysis of T-cells retrieved from blood at day 8/9 following ACT into tumor-bearing mice. Heatmap reflects percentages of the same markers as in a. **c)** FLY-1A TCR T-cells were sorted for CD4^+^ and CD8^+^ using magnetic beads and co-cultured with ROPN1-expressing MM231 cells. IFN-γ and IL-2 production by CD4^+^ and CD8^+^ T-cells upon stimulation is shown (n=2 donors, 2 replicates per donor). Mock T-cells were used as a negative control.

**Extended Data figure 5.**
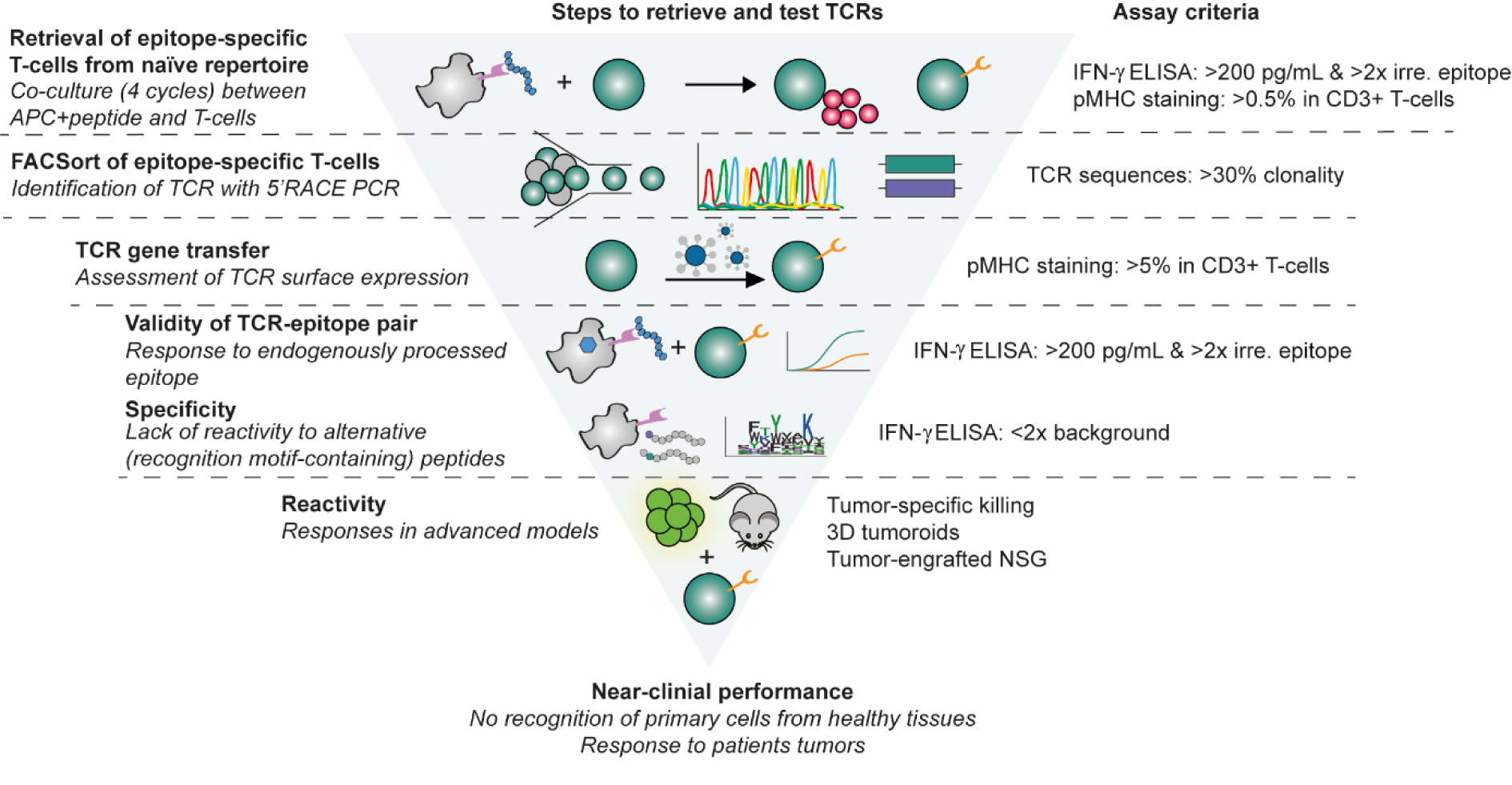
Stepwise approach from retrieval and identification of TCRs to the functional testing of their sensitivity, specificity and reactivity. Cartoon illustrating in 6 steps how ROPN1/B-specific CD8^+^ T-cells are retrieved, the corresponding TCRs identified and tested according to *in vitro, in vivo*, and *ex vivo* assays for sensitivity, specificity and reactivity. Per step, the inclusion criteria that need to be reached for epitope-specific T-cells or TCRs to move to the next step are displayed to the right. Abbreviations in figure: APC, antigen presenting cell; irre. epitope, irrelevant epitope; NSG, NOD scid gamma.

## Extended Data

**Extended Data table 1.**
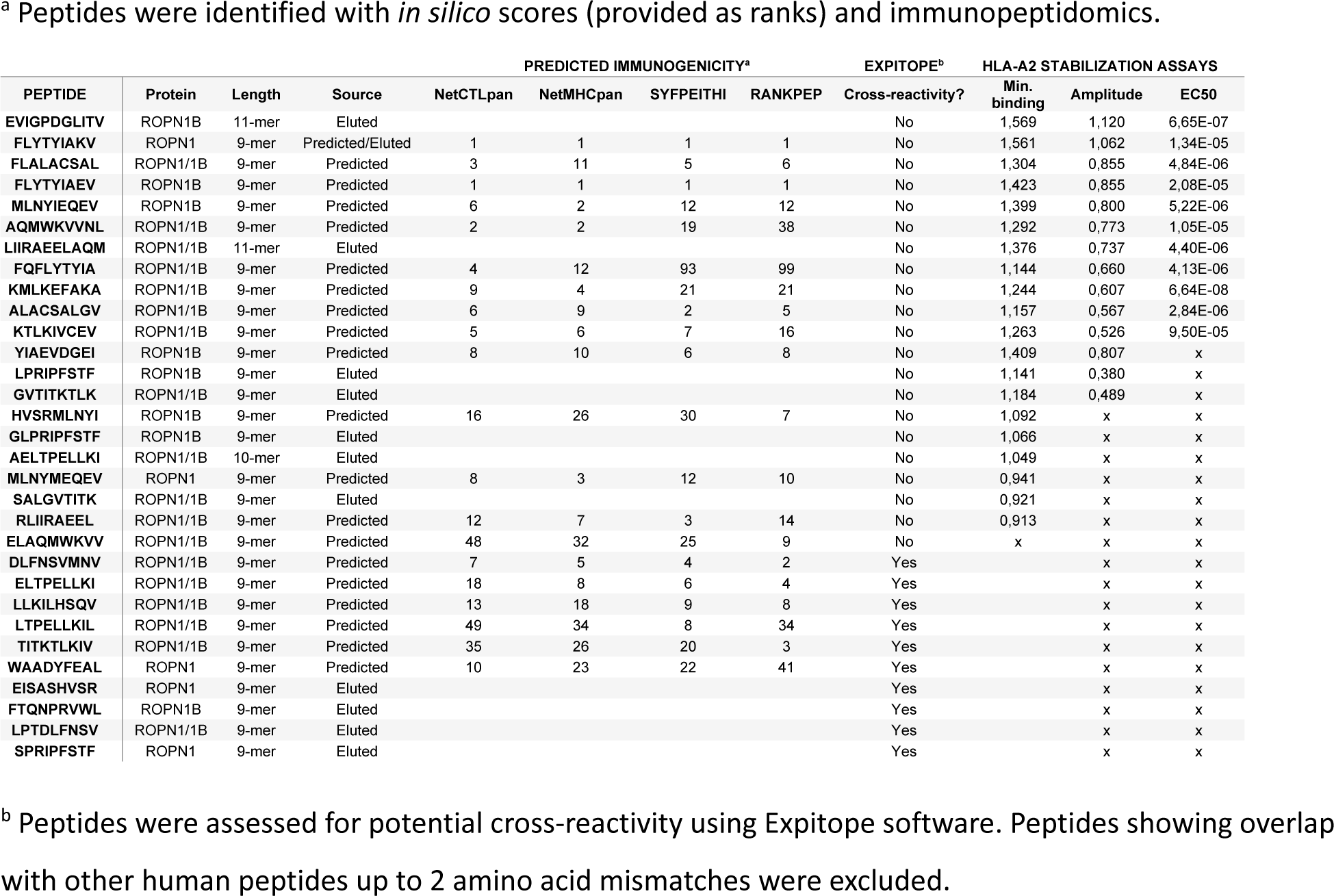
Top-ranked ROPN1/B peptides and their scores for immunogenicity, non-cross reactivity and HLA-A2 binding.

**Extended Data table 2.**
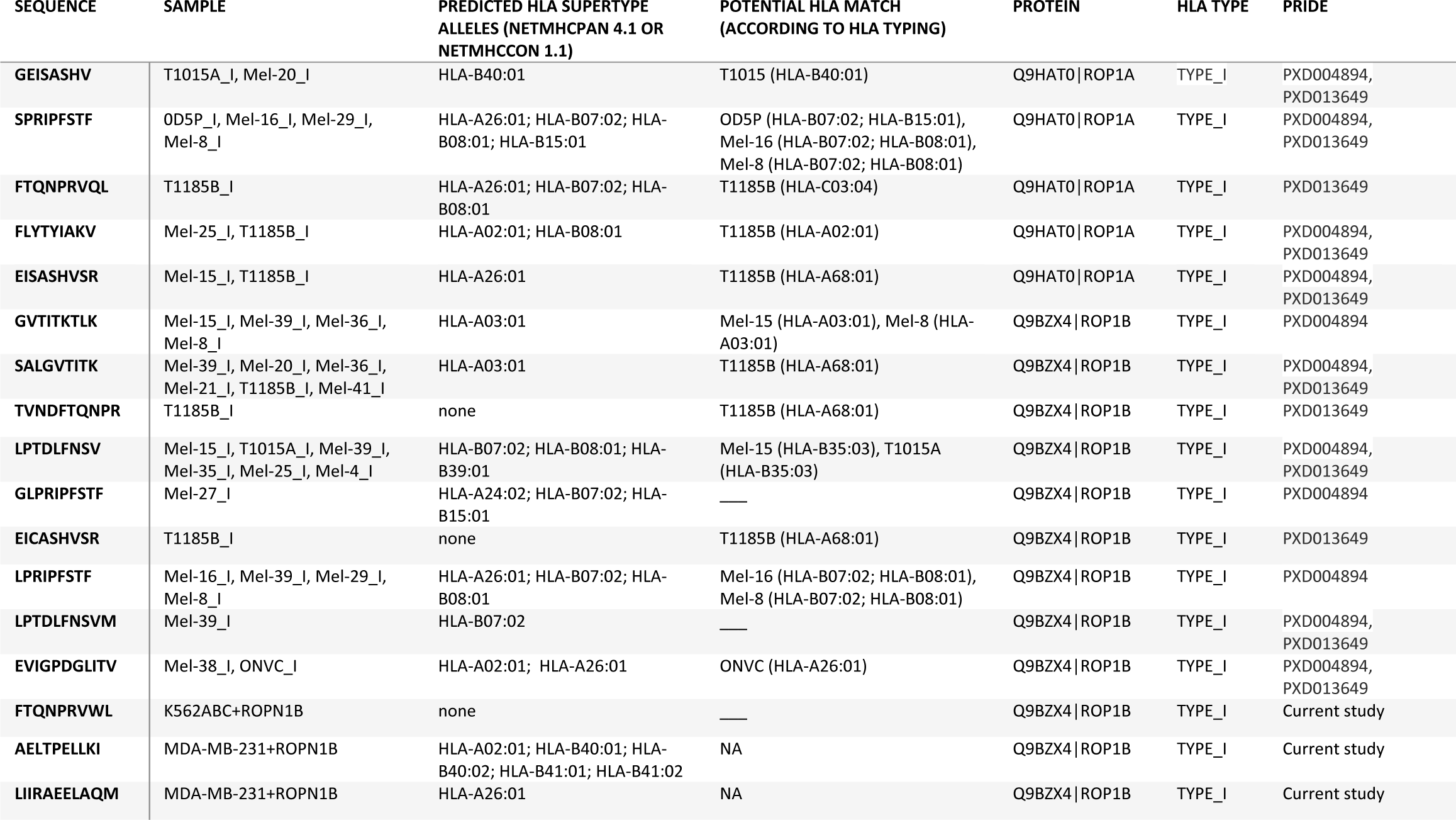
ROPN1/B peptides identified with immunopeptidomics.

**Extended Data table 3.**
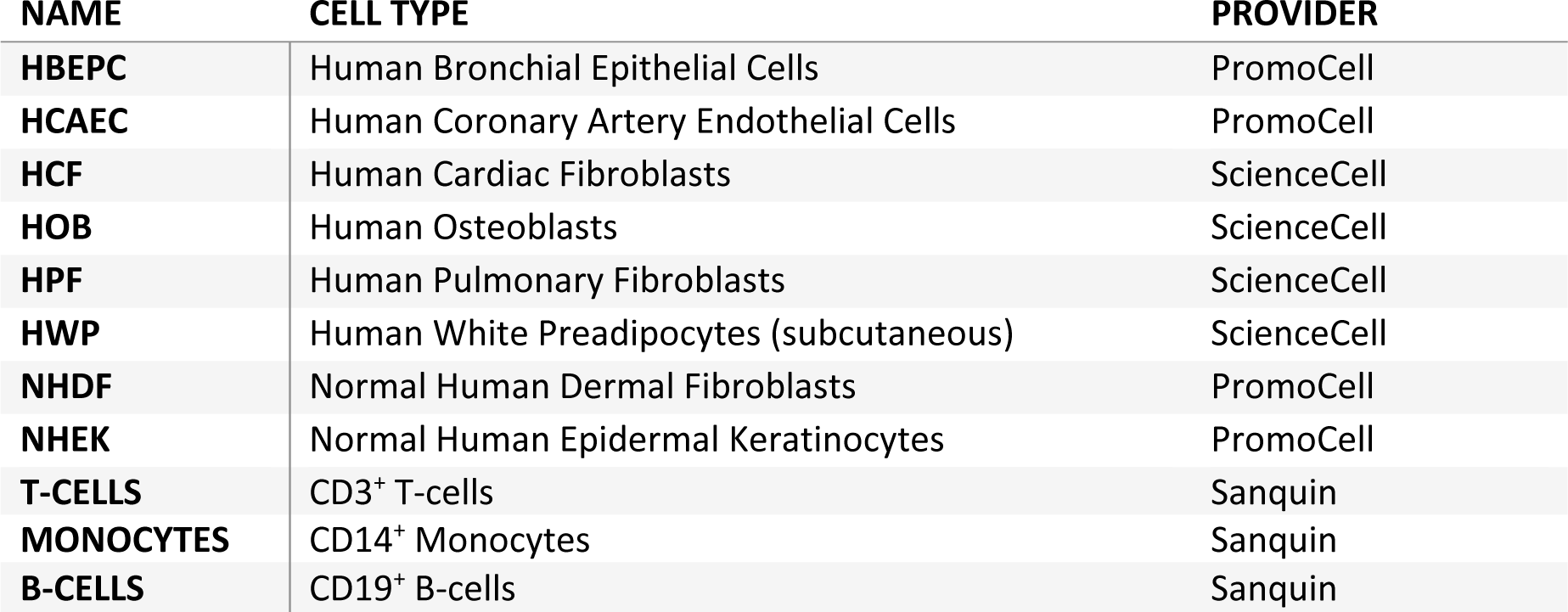
Overview of healthy cell lines.

| **NAME** | **CELL TYPE** | **PROVIDER** |
| --- | --- | --- |
| **HBEPC** | Human Bronchial Epithelial Cells | PromoCell |
| **HCAEC** | Human Coronary Artery Endothelial Cells | PromoCell |
| **HCF** | Human Cardiac Fibroblasts | ScienceCell |
| **HOB** | Human Osteoblasts | ScienceCell |
| **HPF** | Human Pulmonary Fibroblasts | ScienceCell |
| **HWP** | Human White Preadipocytes (subcutaneous) | ScienceCell |
| **NHDF** | Normal Human Dermal Fibroblasts | PromoCell |
| **NHEK** | Normal Human Epidermal Keratinocytes | PromoCell |
| **T-CELLS** | CD3<sup>+</sup> T-cells | Sanquin |
| **MONOCYTES** | CD14<sup>+</sup> Monocytes | Sanquin |
| **B-CELLS** | CD19<sup>+</sup> B-cells | Sanquin |

## List of abbreviations

ACT: Adoptive T-cell therapy
ATP1A3: Sodium/potassium-transporting ATPase subunit alpha-3
BC: Breast cancer
BRCA: Breast cancer
CAR: Chimeric antigen receptor
CGA: Cancer germline antigens
CPCT: Center for Personalized Cancer Treatment
ECM: Extracellular matrix
FAAH1: Fatty-acid amide hydrolase 1
GMP: Good Manufacturing Practice
HER2: Human epidermal growth factor receptor 2
HLA: Human leukocyte antigen
MAGE-A4: MAGE Family Member A4
MHC: Major Histocompatibility Complex
MM231: MDA-MB-231
NBL1: Neuroblastoma suppressor of tumorigenicity 1
NSG: NOD scid gamma
NY-ESO1: New York Esophageal Squamous Cell Carcinoma
ORR: Objective response rate
PARP: Poly ADP-ribose polymerase
PBMC: Peripheral blood mononuclear cells
PD-1: Programmed cell death 1
PD-L1: Programmed cell death ligand 1
PDX: Patient-derived xenograft
pMHC: Peptide-MHC
ROPN1: Ropporin-1
SKCM: Skin cutaneous melanoma
SLC39A8: Metal cation symporter ZIP8
TAP: Transport of associated proteins
TCR: T-cell receptor
TMA: Tissue microarrays
TNBC: Triple-negative breast cancer
TROP2: Trophoblast cell surface antigen 2
UBE2O: (E3-independent) E2 ubiquitin-conjugating enzyme

