## Supplementary methods for "TCR-engineered T-cells directed against Ropporin-1 constitute a safe and effective treatment for triple-negative breast cancer in near-clinical models"

*Generation and culture of cell lines and T-cells*

The parental TNBC cell line MDA-MB-231 (MM231) and its ROPN1/1B-expressing variants were cultured in RPMI medium supplemented with 10% FBS, 200 mM L-glutamine and 1% antibiotics without and with 2 µg/mL puromycin (in case of overexpression varinats). The packaging cell lines 293T and Phoenix-Ampho were cultured in DMEM supplemented with 10% FBS, 200mM L-glutamine, nonessential amino acids, and 1% antibiotics (DMEM complete). The T2 cell line (the LCLxT lymphoblastoid hybrid cell line 0.1743CEM.T2) and the lymphoblastoid cell line BSM were cultured in RPMI medium supplemented with 10% FBS, 200 mM L-glutamine and 1% antibiotics. T-cells were derived from PBMC from healthy human donors (Sanquin, Amsterdam, the Netherlands) by centrifugation via Ficoll-Isopaque (density=1.077 g/cm^3^; Amersham Pharmacia Biotech, Uppsala, Sweden), and following gene transduction (see **Methods**) cultured in RPMI medium supplemented with 25 mM HEPES, 6% human serum (Sanquin, Amsterdam, the Netherlands), 200 mM L-glutamine, 1% antibiotics (T cell medium) and 360 U/mL human rIL-2 (Proleukin; Chiron, Amsterdam, the Netherlands). T-cells were stimulated every 2 weeks with a mixture of irradiated allogeneic feeder cells, as described elsewhere^1^.

*HLA-A2 stabilization assay*

The HLA-A2 stabilization assay was performed using T2 cells as described in^2^ with a slightly adapted protocol. In short, 0.15x10^6^ T2 cells were incubated with (titrated amounts of) peptide for 3h at 37°C/5%CO_2_ in serum-free medium supplemented with 3 µg/mL β2-microglobulin (Sigma). Surface expressed HLA-A2 molecules were measured with flow cytometry using the HLA-A2 mAb BB7.2 (BD Pharmingen, 1:20). To this end, T2 cells were washed, and stained using fluorescently-labeled antibody, incubated for 25 min on ice in the dark, and dissolved in paraformaldehyde (PFA) with 1% FBS. Cells were gated for viability using flow cytometry, and events were acquired on a FACS Celesta (BD) flow cytometer and analyzed using FlowJo software (version 10.7.1, TreeStar, Ashland, OR). In a first screen for HLA-A2 binding, peptides were used at a concentration of 33 µg/mL; in case this resulted in >1.1-fold change (FC) over baseline (T2 cells without peptide), then these peptides entered a second screen and were further titrated from 31 nM to 31 μM.

*Immunopeptidomics and mass spectrometry*

Pellets of 1x10^8^ ROPN1B-expressing K562ABC cells were lysed in PBS buffer containing 0.25% sodium deoxycholate, 0.2 mM iodoacetamide, 1 mM EDTA, 1:200 Protease Inhibitors Mixture (Sigma), 1 mM Phenyl Methyl Sulfonyl Fluoride (PMSF), and 1% octyl-β-D glucopyranoside (Sigma) at 4°C for 1h. Lysates were cleared by 30 min centrifugation at 40,000*g* and supernatants were subjected to sequential immunoaffinity purification of MHC class I with the W6/32 antibody covalently bound to Protein-A Sepharose beads (Invitrogen, Camarillo, CA) using our previously described protocol^3^. To this end, we applied the Waters Positive Pressure-96 Processor and 96-well single-use micro-plates with 3 µm glass fibers and 10 µm polypropylene membranes (Seahorse Bioscience). After loading, the beads in the plates were washed separately with varying concentrations of salts using the processor. Subsequently, the beads were washed twice with 20 mM Tris-HCl, pH8, and eluted with 1% trifluoroacetic acid (TFA; Sigma Aldrich) directly into preconditioned Sep-Pak tC18 100 mg Sorbent 96-well plates (Waters). After an additional wash step with C18 sorbent with 0.1% TFA, MHC class I peptides were eluted with 28% acetonitrile (ACN; Sigma Aldrich) in 0.1% TFA, and recovered peptides were dried using vacuum centrifugation (Concentrator plus, Eppendorf) and stored at −20 °C. Prior to mass spectrometry (MS), peptide samples were re-suspended in ACN/FA (2%/0.1 %), placed in the autosampler of the mass spectrometer, and measured in technical duplicates. MHC-bound peptides were separated by an Easy-nLC 1200 coupled on-line to a Q Exactive HFX mass spectrometer (Thermo Fisher Scientific, Bremen, Germany). A home-made 50 cm long (75 μm inner diameter) column was packed with ReproSil-Pur C18-AQ 1.9 μm resin (kindly provided by dr. Maisch, GmbH) in MeOH. MHC class I peptides were eluted with a linear gradient of 2–28% and 0.1% FA at a flow rate of 250 nl/min over 125 min. MS spectra were acquired from m/z=300-1650 in the Orbitrap with a resolution of 60,000 (m/z=200) and an ion accumulation time of 80 ms. The auto gain control (AGC) was set to 3x10^6^ ions. MS/MS spectra were acquired for the 20 most abundant precursor ions with a resolution of 15,000 (m/z=200), an ion accumulation time of 120 ms and an isolation window of 1.2 m/z. The AGC was set to 2x10^5^ ions, a dynamic exclusion of 20 s and a normalized collision energy of 27. No fragmentation was performed for MHC cl. I peptides in case of assigned precursor ion charge states of four and above. Immunopeptidomics data of K562ABC cells was searched with a PSM false discovery date of 1% with the MaxQuant platform^4^ version 1.6.10.43 against the human proteome UP000005640_9606 reference and a list of 247 frequently observed contaminants. The enzyme specificity was set to non-specific and protein FDR was not set. Peptides with a length between 8 and 25 amino acids were included in our analyses. The initially allowed mass deviation of the precursor ion was set to 6 ppm and the maximum fragment mass deviation was set to 20 ppm. Methionine oxidation and N-terminal acetylation were set as variable modifications.

*Testing reactivity of TCR T-cells towards 3D organoid models*

Organoid models from PDX from breast and skin tumor-derived were established as described^5^. The TNBC PDX BR5010B was expanded in Advanced DMEM/F12 (ThermoFisher) supplemented with 1% Glutamax (ThermoFisher), 1% antibiotics (ThermoFisher), 1% HEPES (Cat#15630-056, ThermoFisher), EGF (5 ng/mL), R-spondin 3 (250 ng/mL), FGF7 (20 ng/mL), FGF10 (20 ng/mL), Heregulin (37.5 ng/mL), SB202190 (500 nM), vitamin B27 (1x), n-Acetyl Cysteine (1.25 mM), Nicotinamide (1M), A83-01 (5 mM), Noggin (100 µg/mL) and Rhoki (10 mM). The melanoma PDX ME12086B was expanded in DMEM/F12 supplemented with EGF (50 ng/mL), ITS (insulin-transferrin-selenium-X supplement) (0.5x), Heparin (2 µg/mL), NEEA (1x), Glutamax (1x), FBS (10x), Rhoki (10 µM) and hydrocortisone (50 nM). On day 0, these 2 PDX organoids were seeded in 3D hydrogels (Proprietary, Crown Bioscience Netherlands B.V.) in 384 well plates (Greiner µClear, Greiner Bio-One B.V.), T-cell medium was added on top and the gel-embedded PDX’es were incubated at 37°C/5%CO_2_ for 48 hours. On day 2, T-cells (i.e., Mock, FLY-1A, FLY-1B or NY-ESO1) were collected and stained with cell tracker (CellTracker™ Green CMFDA Dye, Invitrogen), resuspended in T cell medium and added on top of the 3D hydrogels at different ratios of organoids to T-cells. Cisplatin and Sacituzumab-govitecan, used as drug comp[arators, were added to tumor organoids at 0.1, 1 and 10 µM and 0.1, 1 and 10 nM, respectively. After 48 and 96h, plates were fixed, stained and tumor volumes were analyzed. To this end, cultures were fixed with 4% Formaldehyde (Sigma Aldrich) and simultaneously permeabilized with Triton-X100 (Sigma Aldrich) and stained with rhodamine-phalloidin (Sigma Aldrich) and Hoechst 33258 (Sigma Aldrich) in PBS (Sigma Aldrich) overnight at 4°C, protected from light. After fixation and staining, plates were washed with PBS, sealed with a Greiner SilverSeal (Greiner Bio-One B.V.) and stored at 4°C prior to imaging. Imaging was performed using Molecular Devices ImageXpress Micro XLS with a 4x NIKON objective. For each well, multiple images in the Z-direction were made for both channels, capturing the whole plane in each well. Image analysis was performed using Ominer® software (Crown Bioscience Netherlands B.V.). Individual organoids and immune cells were segmented using detection of Hoechst-stained nuclei, rhodamine-phalloidin-stained cellular F-actin and cell tracker-stained T-cells. These segmentation masks were used to analyze the organoid counts and morphologic profile of organoids per well. Experiments and analyses were performed by personnel at the premises of Crown Bioscience, Netherlands, Leiden.
